# *APOE3*-*R136S* mutation confers resilience against tau pathology via cGAS-STING-IFN inhibition

**DOI:** 10.1101/2024.04.25.591140

**Authors:** Sarah Naguib, Chloe Lopez-Lee, Eileen Ruth Torres, Se-In Lee, Jingjie Zhu, Daphne Zhu, Pearly Ye, Kendra Norman, Mingrui Zhao, Man Ying Wong, Yohannes A. Ambaw, Rodrigo Castaneda, Wei Wang, Tark Patel, Maitreyee Bhagwat, Rada Norinsky, Sue-Ann Mok, Tobias C. Walther, Robert V. Farese, Wenjie Luo, Subhash Sinha, Zhuhao Wu, Li Fan, Shiaoching Gong, Li Gan

## Abstract

The Christchurch mutation (R136S) on the *APOE3* (*E3^S/S^*) gene is associated with attenuation of tau load and cognitive decline despite the presence of a causal *PSEN1* mutation and high levels of amyloid beta pathology in the carrier^1^. However, the specific molecular mechanisms enabling the *E3^S/S^* mutation to mitigate tau-induced neurodegeneration remain unclear. Here, we replaced mouse *ApoE* with wild-type human *E3* or *E3^S/S^* on a tauopathy background. The *R136S* mutation markedly decreased tau load and protected against tau-induced synaptic loss, myelin loss, and reduction in theta and gamma powers. Additionally, the *R136S* mutation reduced interferon response to tau pathology in both mouse and human microglia, suppressing cGAS-STING activation. Treating tauopathy mice carrying wild-type *E3* with a cGAS inhibitor protected against tau-induced synaptic loss and induced similar transcriptomic alterations to those induced by the *R136S* mutation across brain cell types. Thus, suppression of microglial cGAS-STING-IFN pathway plays a central role in mediating the protective effects of *R136S* against tauopathy.

**One-sentence summary:** The R136S mutation on APOE3 enhances resistance to tau-related pathology and toxicity by downregulating the cGAS-STING-IFN signaling pathway.

## Introduction

Alzheimer’s disease (AD) is the most prevalent form of dementia, characterized by pathological amyloid beta (Aβ) plaques and neurofibrillary tangles of hyperphosphorylated tau^2^. Amyloid pathology typically develops early in AD whereas tau pathology occurs later but correlates more accurately with AD progression and cognitive impairment^3–5^. Most AD patients have sporadic late-onset AD (LOAD), in which the apolipoprotein E (*APOE*) gene contains a predominant risk variant^6,7^. APOE is a lipid transporter expressed on astrocytes and microglia and has been shown to affect numerous microglial functions including neuroinflammation and phagocytosis^8–10^. The three major APOE variants, *APOE2*, *APOE3*, and *APOE4*, differ from each other at two amino acid positions, 112 and 158, and modulate AD risk. *E3* is the most common APOE variant and is considered neutral with regards to AD risk, while *E4* alleles elevate AD risk and *E2* reduces AD risk^12,13^. Other variants in APOE have been discovered, such as the *APOE3*-Jacksonville mutation, which reduces amyloid toxicity^11^.

A recent study revealed that a patient carrying the familial *PSEN1* E280A mutation, which causes early onset autosomal dominant AD as early as 30-40 years of age^12^, was protected against cognitive decline until her 70s, a benefit attributed to homozygosity for the *R136S* or Christchurch mutation on *APOE3* background (*E3^S/S^*)^1^. Despite extraordinarily high amyloid burden, the carrier exhibited low tau pathology in her frontal and occipital cortices, typically vulnerable regions in AD^13^, suggesting potent protective functions of the *R136S* mutation against tau-induced neurodegeneration. In mouse models on the risk-enhancing *APOE4* background, the *R136S* mutation was found to protect against tau pathology, neurodegeneration, and neuroinflammation^14^. Additionally, *E3^S/S^* also protected against toxicity induced by inoculation of AD-tau in a transgenic amyloid mouse model^15^. Thus, it appears the protective effects of *R136S* mutation are likely relevant to both *APOE3* and *APOE4* carriers. Furthermore, identifying the mechanisms underlying *R136S-*mediated protection may elucidate potential therapies extending to multiple diseases exhibiting tau pathology.

Our current study aims to dissect the mechanisms underlying the protective function of *E3^S/S^* against tauopathy. We generated human *E3* and *E3^S/S^* knockin models by replacing the expression of mouse *ApoE* with human *E3* and *E3^S/S^* cDNA, followed by crossing with the well-established *P301S* mouse model^16^. We characterized the effects of *E3^S^*^/S^ on tau pathology, synaptic loss, network dysfunction, and demyelination. We performed snRNA-seq to identify cell type-specific transcriptomic changes, and discovered that *ApoE3^S/S^* reduced cGAS-STING-IFN signaling in microglia without affecting disease-associated microglial signatures (DAMs)^17^ or microglial neurodegenerative phenotype (MGnD)^18^. The downregulation of cGAS-STING-IFN signaling in microglia was confirmed in both primary microglia and human iPSC-derived microglial-like cells (iMGLs) following tau treatment. We also determined to what extent the protective function of the *R136S* mutation can be attributed to cGAS inhibition by treating *E3/P301S* mice with a cGAS inhibitor; using snRNA-seq, we compared cGAS inhibitor profiles with transcriptional signatures induced by the *R136S* mutation across several cell types. Our study revealed a central role of cGAS-STING-IFN activation in opposing the *R136S* mutation-induced resilience.

## Results

### *ApoE3^S/S^* mutation reduces tau inclusions and protects against tau-induced loss of synapses

To directly assess the effects of the *R136S* mutation on the *ApoE3* background, we used CRISPR/Cas9 strategies to insert human *ApoE3* (*E3*) cDNA and *ApoE3R136S* cDNA (*E3^S/S^*) into the exon of the mouse *ApoE* locus, resulting in replacement of mouse Apoe with human *E3* or *E3^S/S^* (**Fig. 1A**). PCR confirmed the correct recombination and insertion of human *E3* or *E3^S/S^* cDNA at the *mApoe* locus (**Fig. S1 A–C**). The mouse lines without nonspecific integration for both *E3* and *E3^S/S^* were expanded for subsequent experiments. Sanger sequencing was performed to confirm the accuracy of the replacement (**Fig. S1 B, C**). We then crossed *E3* or *E3^S/S^* mice with PS19 mice, which express the human MAPT transgene with *P301S* mutation (*P301S*), to study effects of *E3^S/S^* on tau pathology in the absence of amyloid, hereafter referred to as *E3/P301S* or *E3^S/S^*/*P301S* mice^16^ (**Fig. 1A**). *E3^S/S^* did not affect APOE levels in the frontal cortex or in plasma in the presence or absence of tau pathology in males (**Fig. 1B, 1C, S1 D**). However, in females, *E3*^S/S^ led to a tau-induced increase in APOE levels in frontal cortex (**Fig. S1 E-F**).

**Figure 1.**
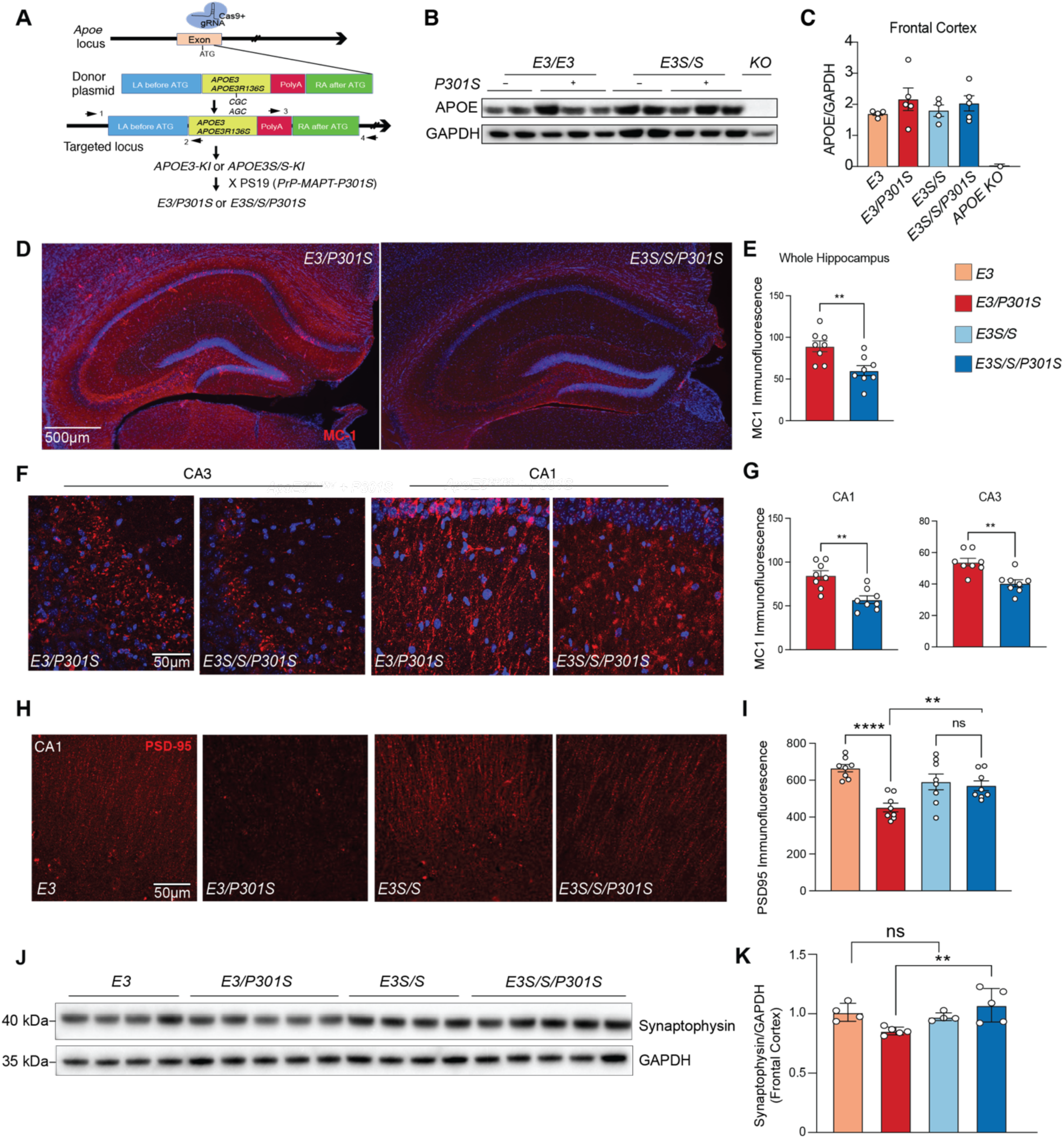
*ApoE3^S/S^* mutation reduces tau inclusions and protects against tau-induced loss of synapses. (A) Schematic illustrating knockin of human *APOE3* or *APOE3R136S* on the mouse *Apoe* locus, and cross with *P301S* mice. (B) Representative western blot for APOE in frontal cortex lysate from all four experimental groups, showing no changes in APOE levels between *E3* genotypes without or with *P301S.* APOE knockout (KO) frontal cortex was used as a control. (C) Quantification of APOE western blot of frontal cortex lysates, showing no differences between groups. Data are reported as mean ± SEM. Data were analyzed by one-way ANOVA with Tukey’s multiple comparison test. (D) Representative immunofluorescence images of whole hippocampus of *E3/P301S* and *E3^S/S^/P301S* mice labeled with MC-1 antibody (red); scale bar, 500um. (E) Quantification of MC-1 immunofluorescence intensities, showing decreased MC-1 throughout *E3^S/S^/P301S* mice. Results are presented as average intensity measures from 3-4 sections per animal with 8 animals/experimental group. Data are reported as mean ± SEM. **p=0.0053. Data were analyzed by unpaired t-test. (F) Representative immunofluorescence images of CA3 and CA1 subregions of hippocampus of *E3/P301S* and *E3^S/S^/P301S* mice respectively labeled with MC-1 (red); scale bar, 50um. (G) Quantification of MC-1 immunofluorescence intensities, showing decreased MC-1 throughout *E3^S/S^/P301S* mice in CA3 and CA1. Results are presented as average intensity measures from 3-4 sections per animal with 8 animals/experimental group. Data are reported as mean ± SEM. **p= 0.0013 and **p for CA1 and CA3 respectively. Data were analyzed by unpaired t-test. (H) Representative immunofluorescence images of CA1 subregion of hippocampus of *E3/P301S* and *E3^S/S^/P301S* mice respectively labeled with PSD-95 (red); scale bar, 50um. (I) Quantification of PSD-95 immunofluorescence intensities, showing decreased PSD-95 throughout *E3^S/S^/P301S* mice in CA1. Results are presented as average intensity measures from 3-4 sections per animal with 8 animals/experimental group. Data are reported as mean ± SEM. ****p<0.0001; *, p=0.0291. n.s, not significant. Data were analyzed by two-way ANOVA with mixed-effects model. (J) Representative western blot for synaptophysin and GAPDH in frontal cortex lysate from all four experimental groups. (K) Quantification of synaptophysin levels normalized to GAPDH levels in western blot of frontal cortex lysates. Data are reported as mean ± SEM. **p = 0.0095 for *E3/P301S* compared to *E3^S/S^/P301S*. Data were analyzed by one-way ANOVA with Tukey’s multiple comparison test.

We next determined the effect of *R136S* on tau pathology in male 9–10-month-old *E3/P301S* and *E3^S/S/^P301S* mice. Immunostaining for MC-1, a conformational specific antibody of aggregated tau^19^, indicated a marked decrease in tau load across the hippocampus in *E3^S/S/^P301S* mice, consistent with previous studies performed on *E4* background (**Fig. 1D-G**)^14^. Using PSD95 immunoreactivity, we measured synaptic integrity and showed that *E3^S/S/^P301S* mice exhibited higher levels of PSD95 immunofluorescence compared with *E3/P301S* mice in the CA1 (**Fig 1. H, I**). Western blot analyses confirmed that tau-induced reduction of synaptophysin in the frontal cortex of *E3/P301S* mice was rescued in the *E3^S/S/^P301S* mice (**Fig. 1J-K**).

### *APOE3 ^S/S^* protects against tau-induced loss of theta and gamma power in hippocampus and cortex

To dissect how *E3^S/S^* affects network activity in tauopathy mice, we performed 16 channel local field potential (LFP) recordings in awake 6–7-month-old male mice exploring an open field chamber (**Fig. 2A**). We did not observe statistically significant differences in animals’ overall speed during open field between any experimental groups (**Fig. 2B**). LFP recording measures amplitudes of oscillations across 0.5–250HZ (**Fig. 2C**). Both theta power (4–8 hz) and gamma power (30 to 100 Hz) are closely linked to key cognitive functions such as attention and memory; previous studies have shown that tau pathology can interfere with neuronal circuits, resulting in deficits in theta and gamma oscillations, long before neurons die^16,20–23^. We quantified theta and gamma power in the somatosensory cortex, the visual cortex, hippocampal CA1, and hippocampal dentate gyrus (DG) regions. The average amplitudes of theta power across brain regions were significantly reduced by tau on *E3* but not *E3^S/S^*background, especially in the DG (**Fig. 2D–F**). While tau pathology appeared to reduce average gamma power regardless of *E3* and *E3^S/S^* background (**Fig. 2G**), it failed to reduce the gamma power in the DG on *E3^S/S^* background (**Fig. 2 H,I**). The protection of the *R136S* mutation against network dysfunction is consistent with delayed cognitive impairment in comparison to others in her kindred^1^.

**Figure 2:**
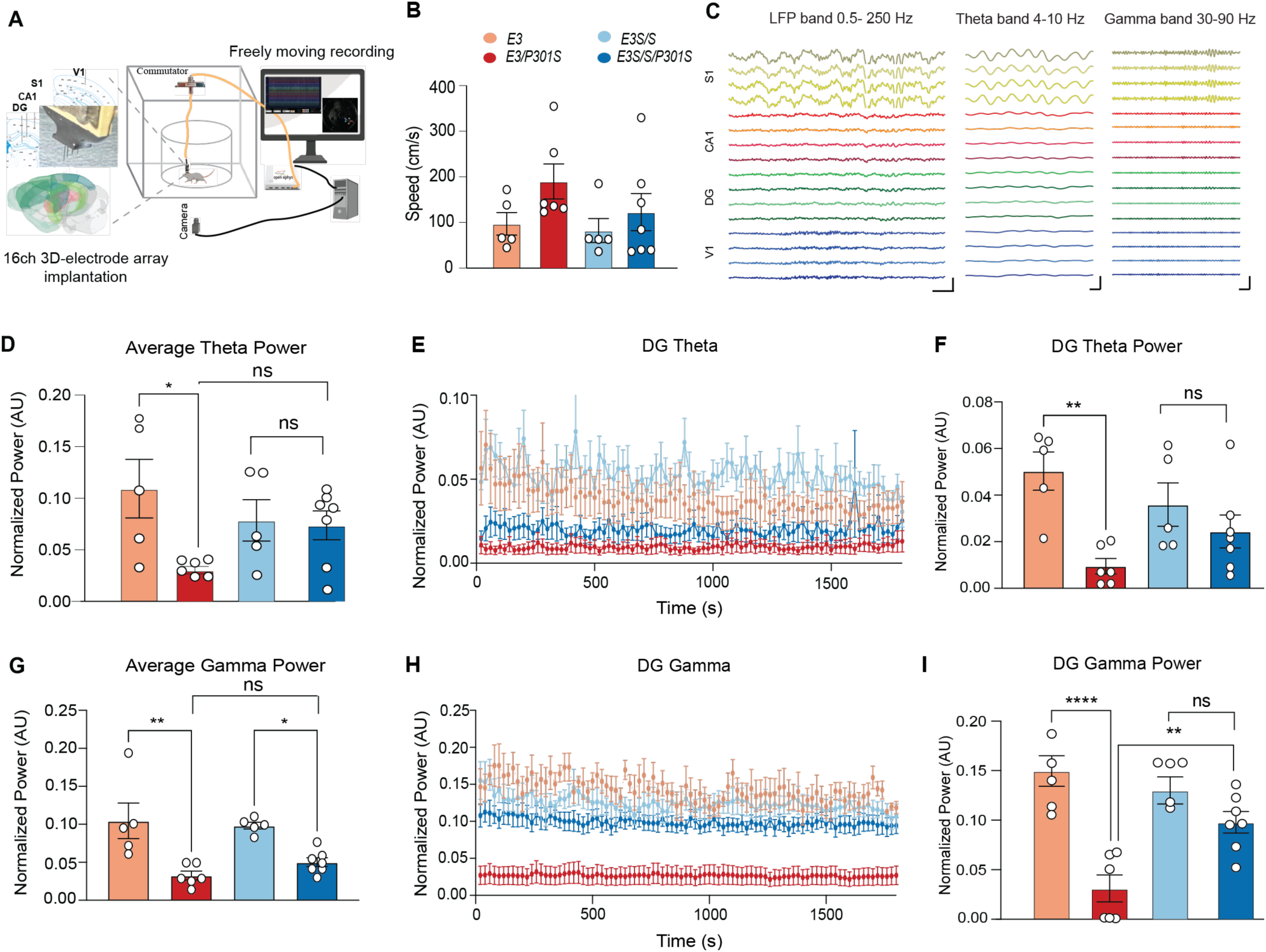
*ApoE3^S/S^* protects against tau-induced loss of theta and gamma power in hippocampus. (A) Schematic of local field potential (LFP) recordings in male 6-7 month old mice in open field chamber. (B) Average speed (cm/s) calculated during LFP recordings from all experimental groups showing no significant difference between groups. N= 5 for *E3*, n=6 for *E3/P301S, n*= 5 for *E3^S/S^*, n= 7 *E3^S/S^/P301S*. Data were analyzed by one-way ANOVA. (C) Representative traces of LFP band, theta band and gamma band respectively for all 16 channels in four brain regions: somatosensory cortex (S1), CA1 of hippocampus, DG of hippocampus and visual cortex (V1). Scale bar represents both amplitude and time (1mV/0.1sec). (D) Quantification of average theta power in all brain regions showing reduction in *E3/P301S* mice in comparison to all other groups. N= 5 for *E3*, n=6 for *E3/P301S, n*= 5 for *E3^S/S^*, n= 7 *E3^S/S^/P301S*. *p=0.0163. Data were analyzed by one-way ANOVA, Tukey’s multiple comparison test. (E) Average theta power from DG in all four experimental groups showing a marked reduction in *E3/P301S* mice in comparison to all other groups (F) Quantification of average theta power in DG showing reduction in *E3/P301S* mice in comparison to all other groups. N= 5 for *E3*, n=6 for *E3/P301S, n*= 5 for *E3^S/S^*, n= 7 *E3^S/S^/P301S*. **, p=0.0042. Data were analyzed byData were analyzed by one-way ANOVA, Tukey’s multiple comparison test.. (G) Quantification of average gamma power in all brain regions across all four groups. N= 5 for *E3*, n=6 for *E3/P301S, n*= 5 for *E3^S/S^*, n= 7 *E3^S/S^/P301S*. **p=0.0016, *p=0.0307. Data were analyzed by one-way ANOVA, Tukey’s multiple comparison test (H) Average gamma power from DG showing a marked reduction in *E3/P301S* mice in comparison to all other groups. (I) Quantification of gamma power in DG showing reduction in *E3/P301S* mice in comparison to all other groups. N= 5 for *E3*, n=6 for *E3/P301S, n*= 5 for *E3^S/S^*, n= 7 *E3^S/S^/P301S*. **p=0.0061, ****p<0.0001. Data were analyzed by one-way ANOVA, Tukey’s multiple comparison test.

Expression of c-Fos, an immediate early-gene marker, is a well-established indicator of mouse brain activity^24^. To examine whether *E3^S/S^* modulates neuronal activity, we performed 3D c-Fos mapping. Eight-month-old male mice from all experimental groups were exposed to a novel open field for 10 minutes, returned to their home cages for 45 minutes, and then their brains were collected for whole-brain c-Fos labeling, clearing, and light-sheet imaging (**Fig. S2A**). Tauopathy increased c-Fos-based brain activity on the E3 background, but this was reversed on the *E3^S/S^* background (**Fig. S2B-C**). Detailed mapping revealed that key regions affected by *E3^S/S^* include the CA1 pyramidal layer (CA1sp), CA1 stratum oriens (CA1so), dorsal subiculum (SUBd), postsubiculum (POST), parasubiculum (PARA), presubiculum (PRE), and entorhinal cortex (ENT) (**Fig. S2D**). These regions, critical for memory retention and retrieval, form the temporoammonic pathway and are highly vulnerable to Aβ and tau pathology^25,26,27^.

### *R136S* mutation induces cell type-specific transcriptomic changes in hippocampi of tauopathy mice

To dissect the cell type-specific effects underlying *R136S*-mediated protection, we then performed single nuclei RNA-seq (snRNA-seq) of hippocampi from male mice in all four genotypes (*E3, E3/P301S, E3^S/S^*, and *E3^S/S/^P301S*) at 9-10 months of age. 88,605 nuclei passed the stringent quality control (QC) (**Fig. S3**). All major cell types were similarly represented across genotypes (**Fig. S3 D, 3A**). The *R136S* mutation in tauopathy mice led to transcriptomic alterations in multiple cell types, including excitatory neurons (EN), inhibitory neurons (IN), astrocytes (AST), oligodendrocytes (OL), microglia (MG), and choroid plexus cells (CHOR) (**Fig. 3B, Table S1**). In the presence of tau, *E3^S/S^* induced the greatest amount of DEGs in CHOR; however, given that only 1,602 CHOR cells were captured in total, we focused primarily on other affected cell types including EN, IN, AST, OL, and MG.

**Figure 3.**
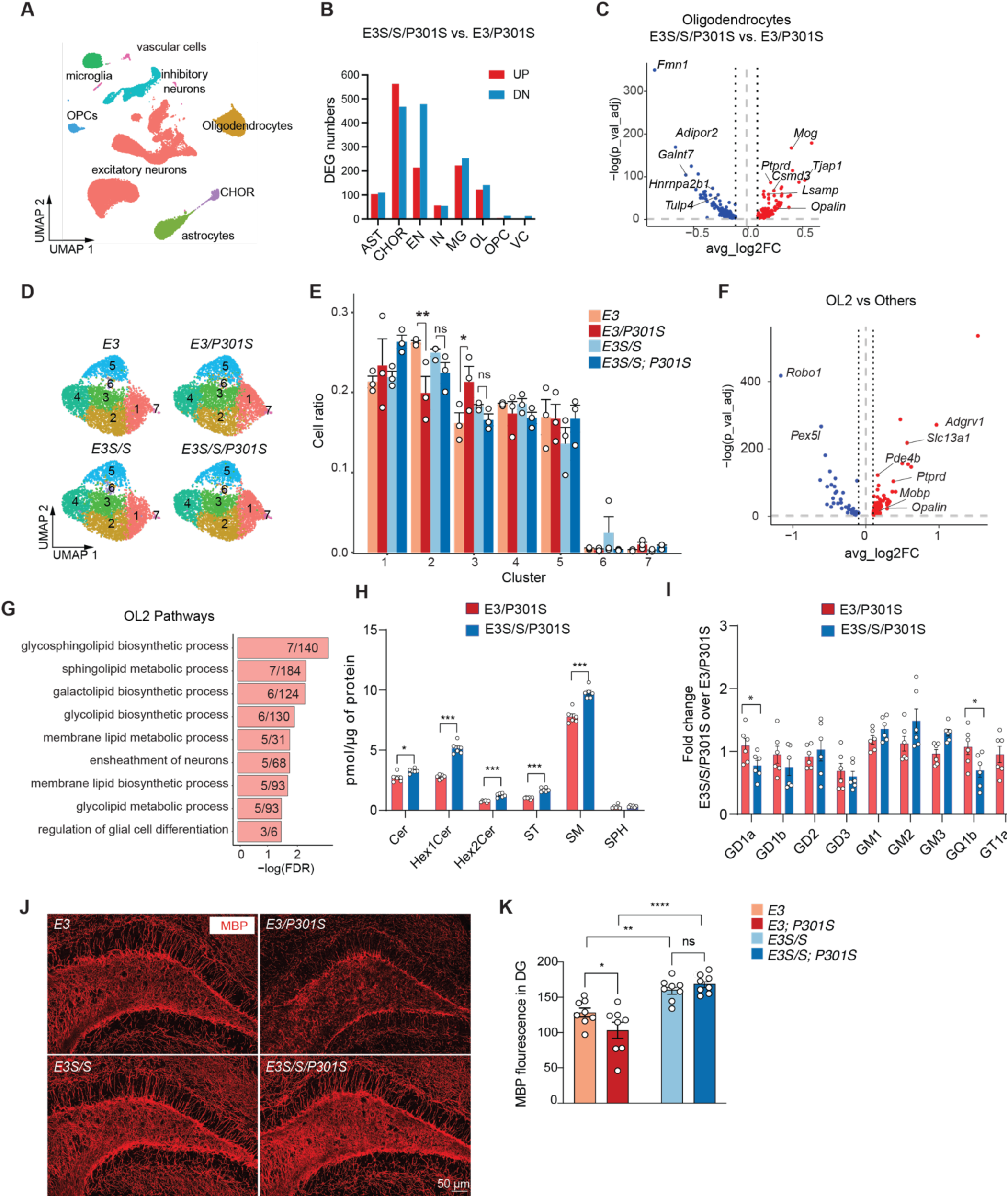
*ApoeE3S/S* induces transcriptomic changes in oligodendrocyte, alters lipid biosynthesis, and protects against tau-induced myelin loss. (A) UMAP of 8 detected cell types using single nuclei RNA-sequencing of the hippocampi from 9–10-month-old *E3* and *E3^S/S^* mice with or without *P301S* tau, n=3 mice in each genotype. (B) Number of significant DEGs between *E3^S/S^* and *E3* in the presence of tau. (C) Volcano plot of DEGs in oligodendrocytes between *E3^S/S^/P301S* and *E3/P301S*. Dashed lines represent log2foldchange threshold of +/– 0.1 and adjusted p value threshold of 0.05. (D) UMAP of oligodendrocyte subclusters split by genotype. (E) Quantification of cell ratios within each subcluster. (F) Volcano plot of oligodendrocyte cluster 2 markers. Dashed lines represent log2foldchange threshold of 0.1 and adjusted p value threshold of +/–0.05. (G) Gene ontology pathway analysis of upregulated oligodendrocytes DEGs from subcluster 2 *E3^S/S^/P301S* vs. *E3/P301S*. (H) Quantification of glycosphingolipids and myelin lipids using LCMS lipidomic analyses, showing that *E3^S/S^/P301S* exhibit higher levels of glycosphingolipids and myelin lipids relative to *E3/P301S.* Lipid classes analyzed included ceramide (Cer), Hexosylceramide (Hex1Cer), Dihexosylceramide (Hex2Cer), Sulfatide (ST), Sphingomyelin (SM), and Sphingosine (SPH). *p < 0.05, ***p < 0.001. Data was analyzed using multiple non-parametric Welch t-tests. (I) Quantification of ganglioside species isolated from the brains of E3S/S/P301S and E3/P301S mice, showing a significant reduction in the fold change of total GD1a and GQ1b in E3S/S/P301S over to E3/P301S. *p < 0.05, with data analyzed using multiple non-parametric Welch t-tests. GM3 (monosialodihexosylganglioside), GM2 (monosialoganglioside), GM1 (monosialotetrahexosylganglioside), GD2 (disialoganglioside), GQ1 (tetrasialotetrahexosylganglioside), and GT1 (trisialotetrahexosylganglioside) are shown. (J) Representative images of immunofluorescence for MBP across each genotype. (K) Quantification of MBP signal across genotypes, n=8 mice per genotype with 3-4 sections per mouse. Data in are reported as mean ± SEM and analyzed via two-way ANOVA multiple comparison. *p<0.05, **p<0.01, ****, p<0.0001

The top upregulated DEGs in ENs were involved in spliceosome assembly and function as well as RNA processing pathways, including *Srsf2*, *Srsf7*, and *Srsf11*, *Rbm25, Rbm39*, *Zranb2*, *Rsrp1*, *Luc7l3* & *Luc7l2* (**Fig. S4A-B**). Using gene set enrichment analysis (GSEA), we found that upregulated genes in ENs also were involved in regulation of membrane potential, inorganic ion transmembrane transport, and RNA processing (*Pnn*, *Pnisr*) (**Fig. S4B**). The primary pathways downregulated in ENs were related to neuronal development and organization (**Fig. S4C).** Although fewer DEGs were observed in interneurons, the top pathways were also associated with RNA processing (**Fig. 3B, S4D).** Comparing the top 50 DEGs between *E3^S/S^/P301S* and *E3/P301S* across excitatory and inhibitory neurons showed that splicing-related genes were shared and predominantly upregulated in both cell types (**Fig. S4E**). Within these shared genes, STRING analysis identified *Rbm25* as a central hub (**Fig. S4F**). To validate the increased expression of *Rbm25* in *E3^S/S^/P301S* on the protein level, we co-immunolabeled for *Rbm25* and NeuN, a neuronal marker, and confirmed that *E3^S/S^/P301S* neurons had higher levels of RBM25 compared to *E3/P301S* neurons (**Fig. S4G-H**).

Astrocytes express high levels of APOE. Immunostaining with an anti-GFAP revealed that tau-induced upregulation of GFAP was not altered by the presence of the *R136S* mutation, suggesting the mutation did not affect tau-induced astrogliosis (**Fig. S4I, J**). SnRNA-seq analyses revealed that the *R136S* mutation led to downregulation of Thioesterase Superfamily Member 4 (*Them4*), which regulates mitochondrial fatty acid metabolism and is involved in apoptotic process (**Table S1, Fig. S4K)**. *Them4* has been shown to regulate PI3K-AKT1 activity, which is involved in toxicity associated with AD risk allele TREM2^28–30^. Another top downregulated gene in astrocytes was metabotropic glutamate receptor 5 (*Grm5*) (**Fig. S4K**). Interestingly, blockade of *Grm5* in human astrocytes reduces pro-inflammatory responses to TNF-α and increases phagocytosis^31^. Among the top upregulated genes are prominent AD risk genes *Apoe* and *Clu*^32^ (**Fig. S4K**). *Clu* overexpression in astrocytes rescued synaptic deficits in an amyloid mouse model^33^. Other upregulated genes are involved in protective functions of astrocytes, such as a glial high-affinity glutamate transporter (*Slc1a2*), which plays a vital role in clearance of glutamate from synaptic clefts, preventing excitotoxicity and thereby protecting neurons^34^ (**Fig. S4K**), consistent with protective effects of the *R136S* mutation against tauopathy.

### *ApoE3^S/S^* induces transcriptomic changes in oligodendrocyte lipid biosynthesis and protects against tau-induced myelin loss

We examined the oligodendrocyte transcriptome in our snRNAseq dataset, performing pseudobulk comparisons between *E3^S/S^P301S* and *E3/P301S* hippocampi (**Fig. 3C**). Top genes upregulated by *E3^S/S^P301S* include myelinating oligodendrocyte markers such as Myelin Oligodendrocyte Glycoprotein (*Mog)* and *Opalin*^35^, a marker of mature oligodendrocytes (**Fig. 3C, Table S1)**. Subclustering analyses revealed that subcluster 2 (OL2) was significantly reduced by tau in *E3* but was preserved in *E3^S/S^* oligodendrocytes (**Fig. 3D-E, Table S2)**. OL2 upregulated several genes associated with myelination including *Pprd*, *Mobp*, and *Adgrv1*^36^ (Fig. 3F). Pathways upregulated by this subcluster were involved in synthesis of several lipid subtypes, including glycosphingolipids and sphingolipids^37^ (**Fig. 3G**). To determine whether this transcriptomic subcluster affected global lipid synthesis, we conducted lipid profiling on *E3^S/S^/P301S* and *E3/P301S* hippocampi (**Fig. 3H**). *E3^S/S^/P301S* brains showed significant increases in total glycosphingolipids (Hex1Cer, Hex2Cer) and myelin (SM, ST) lipids compared with *E3/P301S* brains. In contrast, complex gangliosides GD1a and GT1, which were found to be increased in progranulin deficient model of frontotemporal dementia^38^, showed significant reduction in *E3^S/S^/P301S* (**Fig. 3I**). Changes in other lipid classes were also observed (**Fig. S4 L**). Consistent with the increase in myelin lipid detected in lipidomics, immunostaining with anti-MBP revealed higher levels of myelin in the hippocampus of *E3^S/S^* mice, and a strong protective effect of *E3^S/S^* against tau-induced myelin loss (**Fig. 3J-K)**,

### *ApoE3^S/S^* diminishes tau-induced morphological and IFN responses in microglia

Previous studies showed striking *APOE* isoform-specific effects on innate immune responses including microglia activation^39^, and it has been well-documented that microglia play critical roles in mediating tau toxicity^10,40^. IBA1 immunostaining revealed that tau-induced microgliosis, as measured by IBA1 fluorescence intensity, was not changed with the *E3^S/S^* mutation (**Fig. 4A-B**). Using IMARIS 3D reconstruction to assess microglial morphology, we found that tau increased soma volume in *E3/P301S* microglia, while the *E3^S/S^* mutation prevented this increase (**Fig. 4C-D**). Morphological analysis revealed that the tau-induced decrease in microglial branching was mitigated by *E3^S/S^*, resulting in more branches and a more ramified morphology in *E3^S/S/^P301S* microglia compared to *E3/P301S* microglia (**Fig. 4 C, E**). These findings indicated that the *E3^S/S^* mutation mitigated microglial activation in response to tau pathology.

**Figure 4.**
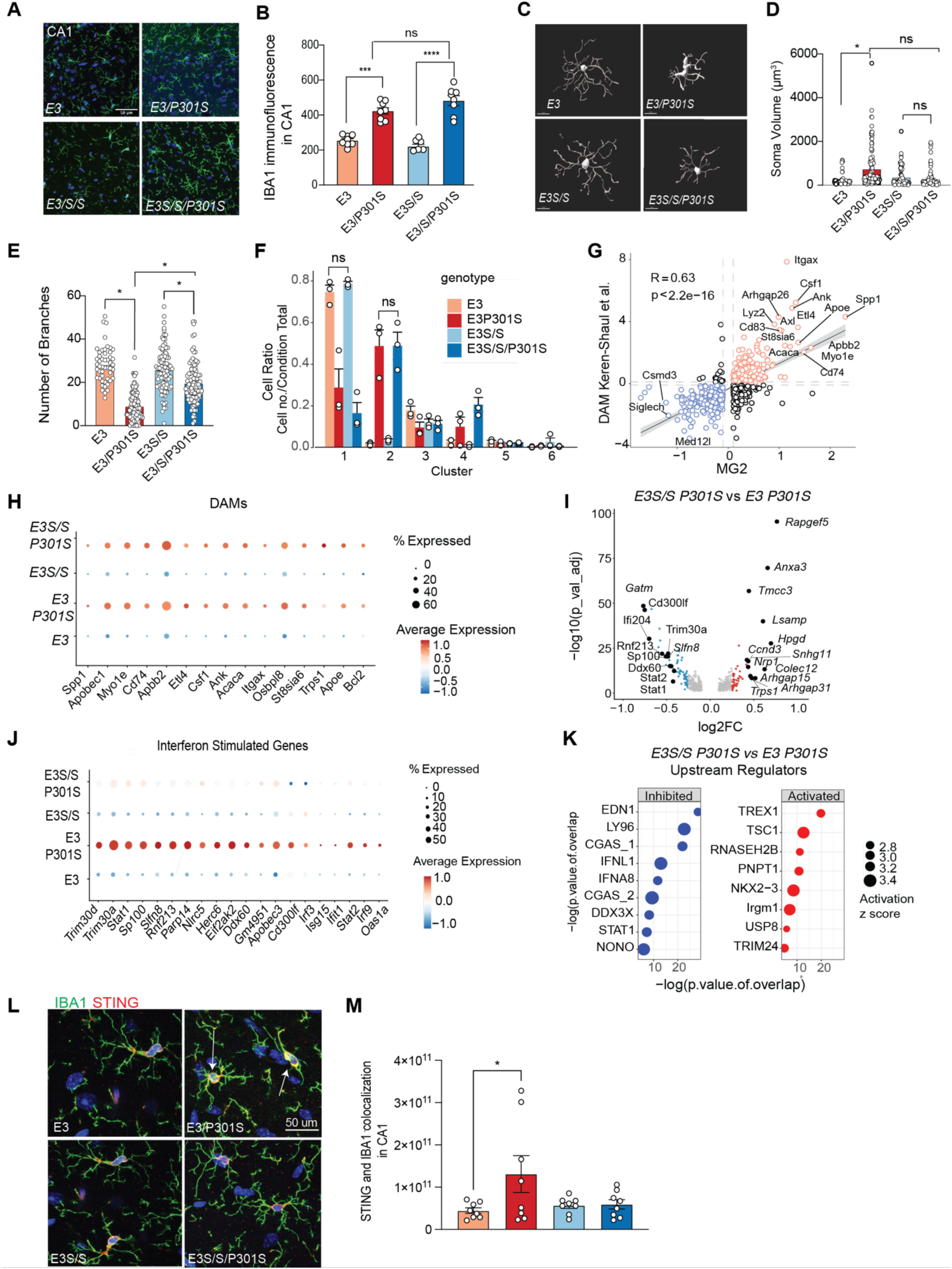
*ApoE^S/S^* rescues tau-induced morphological changes and suppresses IFN responses in microglia. (A) Representative images of IBA1 immunofluorescence from CA1 region of hippocampus in all four experimental groups. (B) Quantification of total IBA1 intensity in CA1 region, showing tau-induced microgliosis not affected by the *E3^S/S^* mutation. Data are reported as mean ± SEM and were analyzed by two-way ANOVA with mixed-effects model, ***p<0.001, ****p<0.0001. (C) Representative IMARIS 3D reconstructed microglia from all experimental groups. (D) Quantification of soma volume size using IMARIS 3D reconstruction in all experimental groups, showing tau-induced increase in soma volume in *E3* microglia that is not present with *R136S* mutation. Each dot represents one microglia, where n=197 microglia for *E3,* n=418 microglia for *E3/P301S,* n=262 for *E3^S/S^,* n=376 microglia for *E3^S/S^/P301S*, from 8 mice/genotype and 4 hippocampal sections/mouse. Data are reported as mean ± SEM and were analyzed by two-way ANOVA with mixed-effects model. (E) Quantification of number of microglial branches using IMARIS 3D reconstruction in all experimental groups, showing tau-induced decrease in branching in *E3* microglia that is significantly prevented in *E3^S/S^* microglia. Each dot represents one microglia, where n=197 microglia for *E3,* n=418 microglia for *E3/P301S,* n=262 for *E3^S/S^,* n=376 microglia for *E3^S/S^/P301S,* from 8 mice /genotype and 4 hippocampal sections/mouse. Data are reported as mean ± SEM and were analyzed by two-way ANOVA with mixed-effects model. (F) Quantification of cell ratios within each microglial subcluster. (G) Correlation between microglia cluster 2 gene expression and published DAM gene signature from Keren Shaul et al. 2017 (H) Dotplot DAM gene expression across four genotypes. (I) Volcano plot of DEGs in excitatory neurons between *E3^S/S^/P301S* and *E3/P301S* thresholded by log2foldchange >0.1 or <-0.1 and adjusted p value thresholded by 0.05. (J) Dotplot of interferon gene expression by genotype. (K) Upstream regulators identified by Ingenuity Pathway Analysis for both up- and downregulated genes by *E3^S/S^* (L) Representative immunofluorescence images of CA1 subregion of hippocampus of *E3, E3/P301S, E3^S/S^, E3^S/S^/P301S* groups co-labeled with Iba1 (green) and STING (red); scale bar, 50um. The arrows indicate colocalization. (M) Quantification of co-localized integrated density of STING in IBA1+ cells. Results are presented as average intensity measures from 3-4 sections per animal with 8 animals/genotype. *p=0.0157 for *E3* compared to *E3/P301S*. Data are reported as mean ± SEM. Data were analyzed by two-way ANOVA with mixed-effects model.

Transcriptomic microglial responses to tau pathology were examined in our snRNA-seq dataset (**Fig. 4F**). Unexpectedly, the cell ratios of homeostatic (MG1) and Disease Associated Microglia (DAM) were similar between *E3^S/S^* and *E3* (**Fig. 4F-H**). Specifically, microglia in *E3/P301S* and *E3^S/S^/P301S* mice exhibited similar loss of microglial subcluster 1 (MG1) enriched with homeostatic markers (**Table S3**), and similar induction of DAM shown by subcluster 2 (MG2) (**Fig. 4F-H, Table S3**). Pseudobulk analysis of DAM genes confirmed similar expression between genotypes (**Fig. 4H**). In contrast, many of the downregulated genes in *E3^S/S^/P301S* microglia were interferon genes, including *Trim30a, Stat1* and *Ifi204* (**Fig. 4I**). Visualizing all genotypes together showed that induction of type-I interferon genes by tau was blunted in *E3^S/S^/P301S* microglia whereas *E3/P301S* counterparts demonstrated a robust type I interferon response to tau (**Fig 4J**). Ingenuity Pathway Analyses confirmed that the top inhibited upstream regulators in *E3^S/S^/P301S* microglia included antiviral activators as cGAS and IFNA, and top activated upstream regulators include those involved in suppressing antiviral response such as TREX1, RNASEH2B (**Fig 4K**). The cGAS-STING pathway was previously shown to play a role in tau-induced neurodegeneration^41^. Immunostaining for STING and IBA1 showed that induction of STING in response to tau occurred solely in E3 microglia (**Fig. 4L-M**), consistent with the notion that the *R136S* mutation dampens cGAS-STING-associated interferon response to tauopathy.

### *ApoE3^S/S^* accelerates tau processing and suppresses tau and Aβ-induced cGAS-STING-IFN response in primary microglia

Our data thus far has pointed to decreased tau load and suppression of microglia cGAS-STING-IFN pathway in *E3^S/S^/P301S* mice. To dissect how the *R136S* mutation affects microglial-mediated tau processing, we isolated primary microglia from brains of *E3* mice and performed a tau pulse-chase assay. Microglia were treated with 0N4R tau fibrils and analyzed at two timepoints: 2 hours post-treatment and 24 hours after extracellular tau supply was halted (**Fig. 5A**). At 2 hours, *E3^S/S^* microglia contained more intracellular tau than E3 microglia; by 24 hours, *E3^S/S^* microglia exhibited less intracellular tau, indicating enhanced tau processing and/or degradation (**Fig. 5B-C**). These findings suggest that *E3^S/S^* microglia show increased tau uptake and clearance, consistent with prior studies^15,42^.

**Figure 5.**
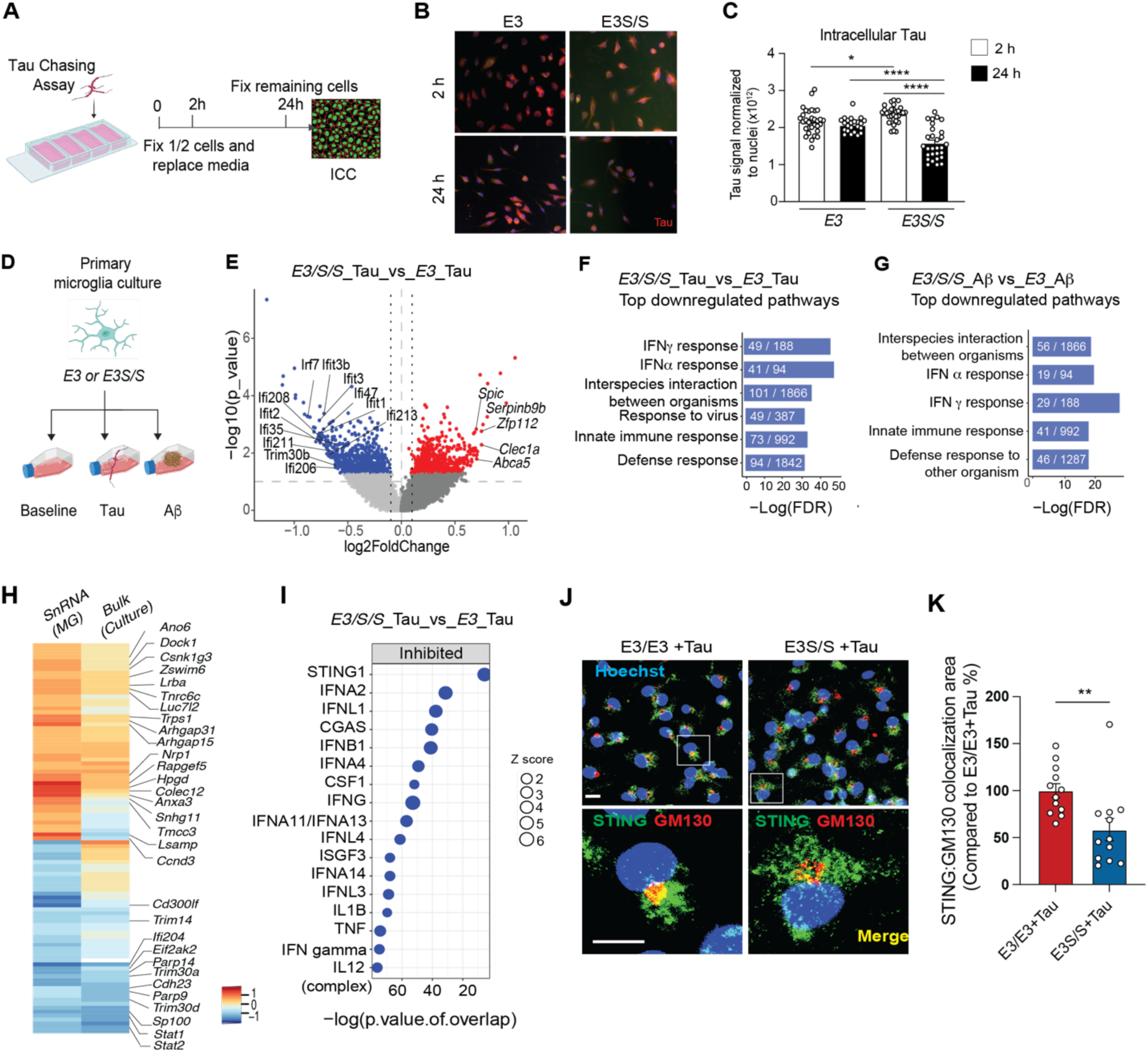
*ApoE3^S/S^* accelerates tau processing and suppresses tau and Aß induced cGAS-STING-IFN response in primary microglia. (A) Schematic for tau chasing assay experiment. (B) Representative immunofluorescence images of *E3* and *E3^S/S^* microglia at 2 hours and 24 hours following tau exposure. Red: anti-tau antibody. (C) Quantification of tau immunolabeling in red fluorescence normalized to number of nuclei/image in *E3* and *E3^S/S^* microglia at 2 hours and 24 hours. Each point on graph represents one well of a chamber slide, with 4-5 images taken/slide. 3 biological replicates were completed (separate microglia dissections) with 8 wells/group each dissection, with n= 122 microglia for *E3* 2 hours, n=90 microglia for *E3* 24 hours, n=117 microglia for *E3^S/S^* 2 hours, and n=114 microglia for *E3^S/S^* 24 hours. Data is reported as mean +/- SEM, and analyzed with two-way ANOVA with mixed effects model. *p=0.0228 for *E3* compared to *E3^S/S^* at 2 hours, ****p<0.0001 for *E3^S/S^* compared to 2 hours compared to 24 hours, and ****p<0.0001 for *E3* compared to *E3^S/S^*at 24 hours. (D) Schematic of treatment plan for primary microglia isolated from either *E3* or *E3^S/S^*pups cultured with basal media, treated with 1 μg/ml 0N4R tau fibrils or 1 μM Aβ42 respectively for 24 hours. (E) Volcano plot of DEGs in primary microglia between *E3^S/S^* + tau and *E3 + tau* thresholded by log2foldchange >0.1 or <-0.1 and p value threshold of 0.05 showing downregulation of interferon genes in bold. (F) Top downregulated pathways in pathways in *E3^S/S^* versus *E3* primary microglia under tau stimulation using GSEA. (G) Top downregulated pathways in pathways in *E3^S/S^* versus *E3* primary microglia under Aβ stimulation using GSEA. (H) Heatmap showing top 50 upregulated and 50 downregulated genes by log fold change in *E3^S/S^* versus *E3* primary microglia under tau treatment compared to those in *E3^S/S^/P301S* mice. (I) Top predicted upstream regulators inhibited by *E3^S/S^* primary microglia compared to *E3* under tau stimulation, including STING and cGAS. (J) Representative confocal images of *E3^S/S^* and *E3* primary microglia labeled with STING (green) and GM130 (red) with merge shown in yellow following 24 hours of tau stimulation. Scale bar represents 20um. (K) Quantification of STING and GM130 colocalization area in *E3^S/S^* and *E3* primary microglia following 24 hours of tau stimulation. Data is reported as mean +/- SEM and analyzed by t-test. N=12 images/condition obtained from 4 wells and two independent experiments. *p<0.01.

We next analyzed the transcriptomic responses of E3S/S microglia to tau or Aβ treatment (**Fig. 5D**). Bulk RNA sequencing revealed downregulation of interferon genes (e.g., *Ifit3, Ifi47, Ifi1, Ifi213, Ifi208, Trim30b, Ifi211, Ifit2, Ifi206*) in tau-treated *E3^S/S^* microglia compared to *E3* counterparts (**Fig. 5E**). GSEA identified IFN-γ response, IFN-α response, and innate immune response as the most downregulated pathways, consistent with findings in microglia of *E3^S/S^/P301S* brains (**Fig. 5F**). Similarly, Aβ-treated *E3^S/S^*microglia downregulated interferon genes (*Ifi211, Ifi204, Cxcl13, Ifi47*), with IFN-γ and IFN-α responses among the top downregulated pathways (**Fig. 5F-G**, **Fig. S5C**). While the downregulated pathways converged, upregulated pathways diverged (**Fig. S5D**). Tau-treated *E3^S/S^*microglia showed enrichment in transcription-related pathways, such as transcriptional regulation, DNA binding, and RNA polymerase II activity, while Aβ-treated *E3^S/S^* microglia upregulated pathways involved the extracellular matrix, plasma membrane, and matrisome (**Fig. S5D**).

To assess whether the most prominent transcriptomic differences between *E3^S/S^* and *E3* microglia were preserved *in vivo* and *in vitro,* we compared the top 50 up and downregulated genes in our snRNA and bulk RNA-seq datasets (**Fig 5H, Table S4**). This analysis revealed shared downregulation of several interferon-related genes in both settings (**Fig. 5H**). Ingenuity Pathway Analysis identified STING1 and cGAS as predicted upstream regulators inhibited in tau-treated *E3^S/S^* microglia (**Fig. 5H-I**). To confirm activation of the cGAS-STING-IFN pathway at the protein level, we performed immunolabeling using a Golgi marker to visualize activated STING in *E3* and *E3^S/S^* microglia treated with tau (**Fig. 5J-K**). Following tau treatment, *E3* microglia showed greater STING/GM130 colocalization compared to *E3^S/S^* microglia (**Fig. 5J-K)**. These findings demonstrate that *E3^S/S^* microglia downregulate the activated cGAS-STING-IFN pathway in response to tau both *in vivo* and *in vitro*.

### *R136S* mutation lowers extracellular tau and suppresses tau-induced cGAS-STING-IFN response in human iPSC-derived microglia

We investigated how *R136S* mutation alters responses to tau in human microglia. *E3* and *E3^S/S^* iPSCs were differentiated into microglia-like cells (iMGLs)^43^ (**Fig. 6A**), confirmed by the expression of microglia markers, IBA1 and TMEM119 (**Fig. 6B**). Both *E3* and *E3^S/S^* iMGLs were treated with recombinant 0N4R tau fibrils, and intracellular tau was assessed after 24 hours. *E3^S/S^* iMGLs exhibited increased intracellular tau compared to E3 iMGLs, while media from *E3^S/S^* cells contained less extracellular tau (**Fig. 6C-E**). The tau-induced interferon response, measured with ELISA revealed that *E3^S/S^* prevented the induction of IP10, an interferon-stimulated factor, seen in *E3* iMGLs (**Fig. 6F**). Immunocytochemistry showed that tau treatment significantly increased levels pSTING and pTBK1 in *E3* iMGLs, but this activation was abolished in *E3^S/S^* iMGLs, indicating that *R136S* suppresses tau-induced activation of the cGAS-STING-IFN pathway (**Fig. 6G-J**). In summary, the *R136S* mutation enhances tau uptake and processing while dampening downstream cGAS-STING-IFN activation in both mouse and human microglia.

**Figure 6.**
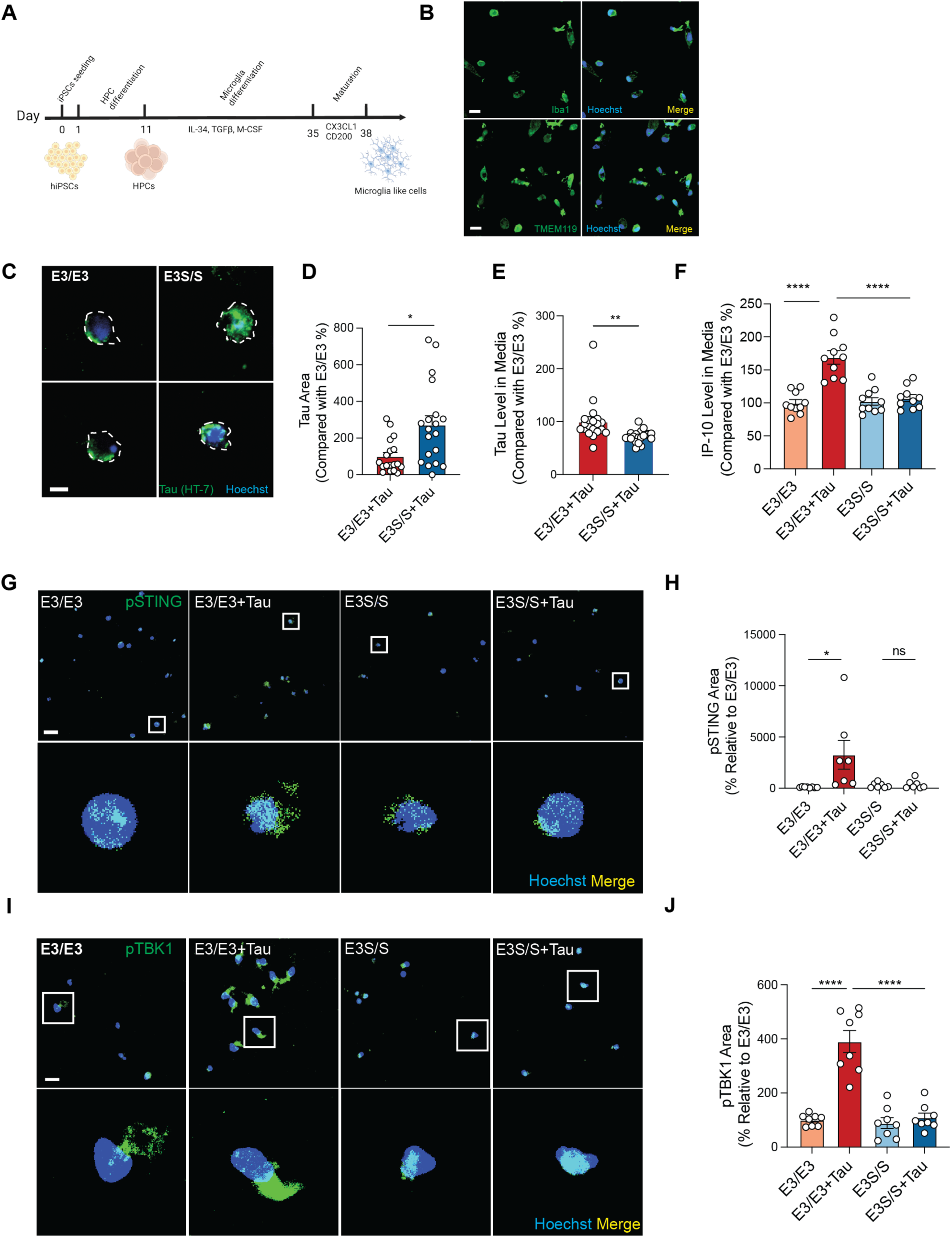
*R136S* mutation suppresses tau-induced cGAS-STING-IFN response in human iPSC-derived microglia. (A) Schematic for human iPSC-derived microglia differentiation. (B) Representative immunofluorescence images of human iPSC-derived microglia expressing classical microglial markers, IBA1 (green, top) and TMEM119 (green, bottom). Scale bar represents 20um. (C) Representative immunofluorescence images of *E3^S/S^* and *E3* human iPSC-derived microglia following 24 hour recombinant tau stimulation, labeled with HT-7 tau antibody (green). Scale bar represents 5um. (D) Quantification of intracellular tau after 24 hours of tau uptake in *E3^S/S^* and *E3* human iPSC-derived microglia. Data is reported as mean +/- SEM and analyzed with two-way ANOVA with mixed-effect model. N=20 represents 20 individual microglia from 4 wells. Experiment was replicated 2 times. Outliers were excluded based on an outlier test. *p<0.05. (E) Quantification of remaining extracellular tau in media after 24 hours of tau uptake in *E3^S/S^*and *E3* human iPSC-derived microglia. Data is reported as mean +/- SEM and analyzed with t-test. N=20 represents media collected from 20 individual wells. The level of tau was normalized with protein concentration. Experiment was replicated 2 times. **p<0.01. (F) IP-10 levels, detected by ELISA, in media after 24 hours of tau stimulation in *E3^S/S^* and *E3* human iPSC-derived microglia. Data is reported as mean +/- SEM and analyzed by two-way ANOVA with mixed-effects model. N=10 represents media collected from 10 individual well. The level of IP-10 was normalized with protein concentration. Experiment was replicated 2 times. ****p<0.0001. (G) Representative immunofluorescence images of pSTING (green) in *E3^S/S^* and *E3* human iPSC-derived microglia at baseline and after 24 hours of tau stimulation. Scale bar represents 20um. (H) Quantification of pSTING (green) in *E3^S/S^* and *E3* human iPSC-derived microglia at baseline and after 24 hours of tau stimulation. Data is reported as mean +/- SEM and analyzed by two-way ANOVA with mixed-effects model. Results are presented as average intensity from three images per well. N=7-8 wells from 2 independent experiments. Outliers were excluded based on ROUT outlier test. *p<0.05. (I) Representative immunofluorescence images of pTBK1 (green) in *E3^S/S^* and *E3* human iPSC-derived microglia at baseline and after 24 hours of tau stimulation. Scale bar represents 20um. (J) Quantification of pTBK1 (green) in *E3^S/S^* and *E3* human iPSC-derived microglia at baseline and after 24 hours of tau stimulation. Data is reported as mean +/- SEM and analyzed by two-way ANOVA with mixed-effects model. Results are presented as average intensity from three images per well. N=8 wells from 2 independent experiments. ****p<0.0001

### cGAS inhibition ameliorates tau spread and mimics transcriptomic and protective effects of *R136S* mutation against synaptic loss induced by tauopathy

Given that cGAS-STING-IFN was among the top downregulated pathways induced by the *R136S* mutation in microglia both *in vivo* and *in vitro*, we investigated whether pharmacological inhibition of the interferon response phenocopies the effects of the *R136S* mutation. Our previous work established the efficacy of a brain permeable cGAS inhibitor (TDI6570, also known as G097, hereafter referred to as cGASi) formulated in chow in protecting against tau-induced deficits in P301S mice^41,44^. To confirm target engagement, P301S mice were treated with cGASi via oral gavage and perfused at 2-or 6-hour post-treatment. Western blot analysis demonstrated that cGASi reduced STING levels and downstream pTBK1 activation in the bulk frontal cortex (**Fig. S6A-D**). Immunohistochemistry further showed a reduction in STING-GM130 colocalization in microglia within 6 hours of treatment, indicating robust inhibition of the cGAS-STING-IFN pathway by cGASi *in vivo* (**Fig. S6E-F).**

The *R136S* mutation reduces tau load in human carriers and in *E3/P301S* mice (**Fig. 1**). To assess the impact of cGAS inhibition on tau spreading, we performed a tau seeding assay in 3–4-month-old P301S mice. One side of the hippocampus was seeded with K18 tau fibrils to induce spreading to the contralateral side. Mice were treated daily with cGASi (15 mg/kg) or vehicle control via oral gavage for 1 month following seeding and then perfused. Immunolabeling revealed significantly reduced MC-1-positive tau spread in cGASi-treated mice compared to controls (**Fig. 7A-C**), consistent with enhanced microglial processing and degradation of tau, similar to the *R136S* mutation.

**Fig 7.**
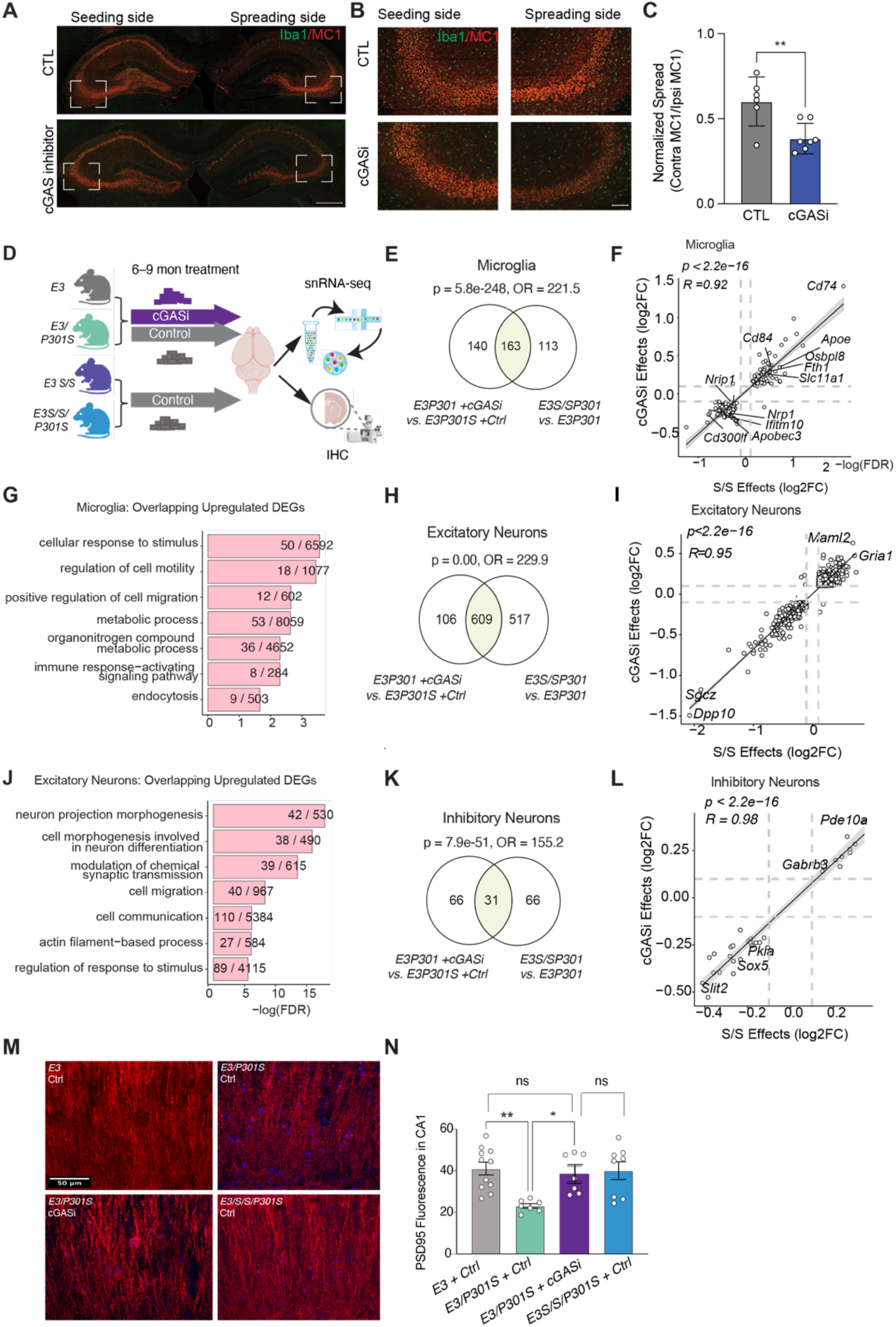
cGAS inhibition ameliorates tau spread and mimics transcriptomic and protective effects of *R136S* mutation against synaptic loss induced by tauopathy. (A) Representative fluorescence images from a tau spreading assay in the hippocampi of P301S mice treated daily with a cGAS inhibitor or control via oral gavage at a dose of 15 mg/kg body weight. Scale bar represents 500um. (B) Higher magnification of CA3 regions of *P301S* hippocampi in tau spreading assay with seeding side on the left and spreading side on the right, MC-1 pathogenic tau labeled in red. Scale bar represents 100um. (C) Quantification of MC-1 positive tau labeling on contralateral side in control and cGAS-i treated *P301S* mice, showing decreased tau spreading following cGAS-i treatment. Results are presented as average intensity measures from 3-4 sections per animal. N=6 mice for CTL; N=7 mice for cGASi. Data are reported as mean ± SEM. **p<0.01 cGASi vs. control. Data were analyzed by unpaired t-test. (D) Schematic plan for 6-month-old *E3* and *E3^S/S^* mice received either a control diet or a cGAS inhibitor diet for 3 months. (E) Venn diagram showing unique effect of cGASi (left) in comparison to effect of *R136S* mutation (right) in microglia as well as overlapping genes between the effects (middle, in green area). (F) Correlation analysis for expression of the overlapping genes in (G). (G) Top upregulated pathways for overlapping genes in (G). (H) Venn diagram showing unique effect of cGASi (left) in comparison to effect of *R136S* mutation (right) in excitatory neurons as well as overlapping genes between the effects (middle, highlighted in green). (I) Correlation analysis for expression of the overlapping genes in (J). (J) Top upregulated pathways for overlapping genes in (J). (K) Venn diagram showing unique effect of cGASi (left) in comparison to effect of *R136S* mutation (right) in inhibitory neurons as well as overlapping genes between the effects (middle, highlighted in green). (L) Correlation of gene expression for overlapping genes in inhibitory neurons in (M). (M) Representative immunofluorescence images of PSD95 staining in CA1 subregion of hippocampus of *E3* mice treated with control diet*, E3P301S* mice treated with control diet*, E3P301S* mice treated with cGAS inhibitor diet, and *E3^S/S^P301S* mice treated with control diet. (N) Quantification of PSD95 immunofluorescence intensity for the corresponding conditions in (M), n=11 animals for *E3/control,* n=7 animals for *E3/P301S/control,* n=8 animals for *E3/P301S/cGASi,* n=8 animals for *E3^S/S^/P301S/control* with 3-4 sections per mouse. Data are reported as mean ± SEM. Data were analyzed by one-way ANOVA with Tukey’s multiple comparison, *p=0.025; **p=0.0028.

We then assessed to what extent cGAS inhibition mimics the protective effects of the *R136S* mutation post the disease onset by treating 6-month-old *E3/P301S* mice—after tau-induced network dysfunction onset—with a control or cGASi diet for 3 months, followed by snRNA-seq and immunohistochemistry (**Fig. 7D**). A total of 205,400 nuclei were analyzed after stringent quality control, with all major cell types similarly represented across genotypes (**Fig. S7**). We compared cell type-specific transcriptomic changes induced by cGASi in *E3/P301S* mice to those observed in *E3^S/S^* mice, both in the presence of tau pathology. In microglia, cGASi treatment induced 163 differentially expressed genes (DEGs) overlapping with those regulated by the *R136S* mutation, with highly correlated expression patterns (R=0.92, **Fig. 7E-G, Table S5**). Pathway analysis of these overlapping DEGs revealed that both cGASi and the *R136S* mutation upregulated pathways related to cell motility, endocytosis, migration, and metabolic processes (**Fig. 7G**). Similarly, striking overlaps and strong correlations in expression patterns were observed in excitatory neurons (ENs; R=0.95, **Fig. 7H-J, Table S5**) and inhibitory neurons (INs; R=0.98, **Fig. 7K-L, S6E, Table S5**). In ENs, shared pathways included cell morphogenesis, synaptic transmission, cell migration and communication, and regulation of response to stimuli (**Fig. 7J**). INs were not included in pathway analysis due to a low number of upregulated DEGs (<30). These findings strongly support that cGAS inhibition transcriptomically mimics the *R136S* mutation in both neurons and microglia. Furthermore, cGASi treatment ameliorated PSD95 loss in E3/P301S tauopathy mice, restoring levels to those observed in *E3^S/S^/P301S* mice (**Fig. 7M-N**). However, cGASi did not alter levels of demyelination in E3/P301S mice as determined with MBP immunofluorescence, suggesting that the protection against demyelination seen in *E3^S/S^* mice may involve a cGAS-independent mechanism, potentially regulated by Tlr7^45^ (**Fig. S7 H, I**). In summary, cGAS inhibition reduced tau spread, and treatment initiated after disease onset recapitulates some key aspects of the *R136S* mutation-mediated resilience to tau toxicity, including transcriptomic changes in neurons and microglia and preservation of synaptic integrity.

## DISCUSSION

Our current study combines functional assays, transcriptomic analyses, and pharmacological intervention to shed light on mechanisms enabling *E3^S/S^* to protect against tauopathy. We established new *APOE3* and *APOE3^S/S^* knock-in mice and demonstrated that the *E3^S/S^* mutation protects against tau-induced declines in theta and gamma oscillations. *E3^S/S^* mice exhibited decreased tau load and tau-induced synaptic loss. The *E3^S/S^* mutation affects multiple cell types, and specifically downregulates cGAS/STING-IFN pathway in microglia. In primary microglia, the *E3^S/S^* mutation enhances tau processing, and suppressed tau-induced cGAS/STING-IFN pathway. This finding was also recapitulated in human iPSC-derived microglia. Remarkably, treating *E3/P301S* mice with a cGAS inhibitor after the onset of network dysfunction phenocopied the *E3^S/S^* mutation on microglial and neuronal transcriptomes, and protected against tau-induced synaptic loss.

In our current model, we replaced the expression of the mouse *Apoe* gene with human *E3* and *E3^S/S^* cDNA. Consistent with observation in an earlier study, in which genomic *APOE3* was used to replace mouse *Apoe* alleles, the *R136S* mutation did not affect brain APOE3 levels in males^15^. Interestingly, we observed an increase in APOE in female *E3^S/S^/P301S* cortex. However, unlike Chen et al^15^, which reported elevated plasma APOE levels in male *E3^S/S^* mice, we observed similar APOE levels in *E3* and *E3^S/S^* mice in male plasma. APOE levels were not reported to be affected by *R136S* mutation on the APOE4 background in the Nelson et al study^14^. Exactly how the *R136S* mutation affects APOE3 metabolism in brains vs. plasma and how sex could modify the effects remains unclear. Nevertheless, the comparable APOE levels in the brain and plasma of our male *E3* and *E3^S^*^/S^ mice allow us to investigate the specific effects induced by *R136S* mutation without the confounds caused by different levels of APOE isoforms.

The *E3^S/S^* patient had lower tau load in comparison to other individuals with the same PSEN1 mutation, and very high amyloid pathology^1^. We showed that *E3^S/S^/P301S* mice demonstrated less tau pathology across the hippocampus, like previously published observations on the *APOE4* background^14^. On the *E3* background, previous studies elucidated robust protective effects in an amyloid model induced by tau aggregates isolated from AD brains (AD-tau)^15^. Thus, while the protective effects of the *R136S* mutation were discovered on an *E3* carrier with amyloid pathology, it is likely that the protective mechanisms of the *R136S* mutation are broadly applicable, not only to E3 and E4 carriers, but also to FTD-tau and primary tauopathies.

How does *R136S* mutation lower tau load? Previous studies point to neuronal or microglial mechanisms. So far, all *E3^S/S^* knockin models are whole-body knockin; conditional knockin models are needed to further solidify cell type-specific mechanisms. In primary microglia, we observed that *E3^S/S^* microglia have more intracellular tau than *E3* microglia at 2 hours, indicating that *E3^S/S^* microglia are more efficient at tau uptake at early timepoints than *E3* microglia. After external tau was removed at 2 hours, we found that the *E3^S/S^*microglia had less intracellular tau than *E3* microglia—indicating that the *R136S* mutation results in more efficient tau processing. Our *E3^S/S^* human-derived iPSC microglia also take up tau more efficiently than *E3* microglia: after 24 hours of constant tau stimulation, we found increased intracellular tau in conjunction with less extracellular tau. Our data aligns with recent studies using *E3^S/S^* microglia co-cultured with either PSEN1 neurons or Aβ-42 in which the *R136S* mutation increases phagocytic ability and reduces phosphorylated tau load to protect neurons^42^. While the exact mechanism by which the *R136S* mutation reduces tau load is yet to be determined, our data indicates a role for the cGAS-STING-IFN pathway. Treatment of cGAS inhibitor significantly reduced tau spread, a process in which microglia are critically involved. Additional studies are needed to determine if dampening the cGAS-STING-IFN pathway can also increase efficiency of amyloid processing, as we also observed a reduction in interferon pathways following amyloid treatment in primary microglia.

Using LFP recording of network activities, we showed that *E3^S/S^* mice are protected against tau-induced deficits in theta and gamma oscillations in multiple brain regions at 6-7 months of age, before tau-induced neurodegeneration at 9-months of age, supporting the notion that network dysfunction represents early pathogenetic mechanisms in AD^46^. Previous studies have shown that amyloid pathology induces abnormal theta and gamma oscillations, likely due to deficient inhibitory interneuron activity, which we did not examine in this study^20,47,48^. The revelation that frontotemporal dementia (FTD) tau pathology led to similar network dysfunction suggests converging network dysfunction in AD and primary tauopathies^49–51^. Indeed, gamma frequency modulation was found to be beneficial in models with either amyloid and/or tau pathology^52^.

Our snRNA-seq analyses of excitatory and inhibitory neurons revealed striking upregulation of RNA-splicing pathways by the *R136S* mutation. Whether it is the consequence or causative of reduced tau load remains an open question that must be followed up in future studies. Neurogenesis is among the downregulated pathways in *E3^S/S^/P301S* hippocampus. Neurogenesis is known to be elevated in response to hyperexcitability^53,54^, which was observed using cFOS activity mapping in *E3/P301S* mice. The downregulation of c-FOS activity by the *R136S* mutation could help explain the reduction in neurogenesis. The *R136S* mutation also exerted a distinct effect on the transcriptomes of astrocytes, including upregulation of protective astrocytic genes, such as EAAT2, the main glutamate transporter that exerts protective function against excitotoxity. Its expression diminishes with age^55^ and in postmortem brains of patients with sporadic AD correlates with signs of neuronal death^56^. Given that agonist of EAAT2 has been found to be neuroprotective^57^, confirming and dissecting how the *R136S* mutation enhances expression of EAAT2 may be of therapeutic potential.

AD patients exhibit increased myelin loss, and higher levels of myelin are associated with protection against tau pathology in AD^43,44^. Our findings suggest that the R136S mutation alters oligodendrocyte transcriptomes, providing protection against tau-induced myelin loss, as evidenced by both immunohistochemical and lipidomic analyses. Notably, APOE4 alone has also been linked to demyelination, even in the absence of tau pathology^45^. In addition, transcriptomic changes were observed in a specific oligodendrocyte subpopulation on an E4 background with the *R136S* mutation^14^. Our lipidomics study reveal elevated levels of Hex1Cer, Hex2Cer, and SM (sphingomyelin) in *E3^S/S/^P301S*, suggesting enhanced glycosphingolipid and sphingolipid synthesis. This elevation likely contributes to increased myelination, Interestingly, we also observed a reduction in complex glycosphingolipids, GD1a and GT1b gangliosides and GALNT7, a gene involved in ganglioside synthesis, in *E3^S/S/^P301*S. These gangliosides are known to form toxic aggregates commonly associated with neurodegenerative diseases, such as frontotemporal dementia (FTD)^38^ and Gaucher’s disease^58^. Dissecting how *R136S* mutation enhances glycosphingolipid and sphingolipid synthesis while mitigating the accumulation of neurotoxic gangliosides could shed new light into its protective mechanisms.

Maladaptive microglia play a central role in pathological tau spread and tau toxicity^40,59^. While the exact toxic pathways remain elusive, several recent studies provide compelling evidence supporting interferon pathways to be particularly damaging, especially in the presence of tau pathology. In tauopathy mouse models, our previous studies suggest that the source of IFN signaling could be partially attributed to hyperactive cGAS-STING in tauopathy mice and the human AD brain^41^. *Cgas* deletion mitigated tauopathy-induced microglial IFN-I and protected against synapse loss, synaptic plasticity, and cognitive deficits^41^. Moreover, *APOE4* greatly exacerbates tau-induced neurodegeneration, partially attributed to a critical immune hub that involves IFN-driven microglial activation and interaction with cytotoxic T cells^60,61^. Previous findings showed that R136S mutation enhances microglial response around amyloid plaques on *E3* background^15^, but downregulates disease-associated gene profiles on the *E4* background^14^. Our current findings reveal that the resilient *E3^S/S^* allele selectively suppresses tau-induced cGAS-STING-IFN signaling in microglia while maintaining DAM responses, providing a protective mechanism that mitigates tau pathology and preserves cognitive and synaptic functions by balancing immune activation and resilience pathways.

Upon tau stimulation, primary microglia from *E3^S/S^*mice exhibit reduced expression of interferon-stimulated genes, indicating that the *R136S* mutation downregulates microglial cGAS-STING-IFN pathway in both *in vivo* and *in vitro*. To determine whether this attenuation is cell-autonomous, future studies should employ microglia-specific knock-in models of the *R136S* mutation to evaluate its sufficiency in providing neuroprotection and suppressing interferon gene expression via the cGAS-STING pathway. Notably, the dampening effect of the *E3^S/S^* mutation on tau-induced cGAS-STING activation was replicated in human iPSC-derived microglia, suggesting that this suppression of the interferon response to tau is conserved in human. Previous research has demonstrated that the *R136S* mutation impairs APOE3’s interaction with heparan sulfate proteoglycans (HSPGs). Marino et al. developed an APOE3 Christchurch-inspired antibody to disrupt human APOE4-HSPG interactions, resulting in reduced tau pathology in the retinas and brains of *P301S* mice^62^. In macrophages, HSPGs modulate inflammation through interferon-beta (IFN-β); specifically, HSPGs sequester IFN-β, and the degree of proteoglycan sulfation can influence inflammatory responses^63^. Further investigation is required to determine whether similar mechanisms operate in microglia and how the impaired interaction between *R136S* and HSPGs leads to suppression of the interferon response following tau stimulation.

Pharmacological inhibition of cGAS in *E3/P301S* mice after the onset of network dysfunction was employed to compare the protective effects of chronic inhibition with those conferred by the *R136S* mutation on the *E3* background. Consistent with previous studies in tauopathy mice expressing mouse Apoe, cGAS inhibition mitigated tau-induced synaptic loss. The significant overlap and striking correlation of transcriptomic alterations across multiple cell types suggest that the resilience observed in the human *R136S* carrier can be partially replicated through pharmacological cGAS inhibition even after the disease onset. cGAS inhibition, however, did not prevent myelin loss, as observed with the *R136S* variant, indicating that the R136S mutation may confer additional cGAS-independent protective effects, or that earlier intervention is required, or both. Future studies elucidating the relationship between APOE isoforms and cGAS activation may prove useful in unraveling shared mechanisms and therapeutic potential of cGAS inhibitors in AD and related dementia. Collectively, this study elucidates the impact of the *R136S* mutation on tau-induced network activity and function, reveals extensive transcriptomic effects on neurons and glial cells, and identifies a potential therapeutic pathway for intervention.

## ACKNOWLEDGEMENTS

The study was supported by: NIH R01AG072758 (to LG), R01AG074541 (to SCS, LG), 1R01AG079291-01A1 (to LG), R01AG079557-01 (to LG), 1F32AG085960-01 (to SN), JPB Foundation (to LG), Rainwater Charitable Foundation (to LG), Cure Alzheimer Fund (to SCS, LG), Gilliam Fellowship from Howard Hughes Institute (to CLL). We would like to thank Ravi Kumar Nagiri for his assistance in producing TDI-6570 chow for use in this study. We would like to thank Dr. Yueming Li’s group for providing Aβ monomers for this study.

## AUTHOR CONTRIBUTIONS

Conceptualization: LG, SN, SG, ERT

Methodology/Investigation: SN, CLL, ERT, SIL, LF, JZ, DZ, PY, KN, MZ, MYW, YA, WW, TP, MB, RN, SM, TW, RF, WL, SS, ZW

Visualization: SN, CLL, SIL, MZ, LF, RC, ZW

Funding Acquisition: LG, SS, SN, CLL

Writing – original draft: SN, CLL, LG

Writing – reviewing and editing: SN, CLL, LG

## MATERIALS/METHODS

### Generating Mouse Lines

To introduce a human *APOE3 and APOE3Christchurch (APOE3-R136S, E3S/S)* cDNA into mouse *Apoe* genetic locus, a donor plasmid was designed and made. The human *APOE3* or *APOE3S/S* cDNA and a poly A tail were flanked by left and right homology arms which were designed to insert human *APOE3* or *APOE3S/S* right before the start codon of mouse *Apoe* gene. Upon recombination, the mouse *Apoe* promoter and regulatory elements would drive expression of the inserted human *APOE3* or *APOE3S/S* cDNA, whereas the expression of mouse *Apoe* gene would be inactivated. A single guide RNA (sgRNA) was selected. RNP containing sgRNA and Cas9 protein was prepared. Pronuclear injection of RNP and the donor plasmid was performed at the Rockefeller University transgenic core facility. Genomic DNA from founders was isolated from tail lysate. To screen the specific knockin mice, two PCR amplifications were performed with primers flanking the outside of the homology arms and on the human APOE3 transgene. The PCR products were sequenced to validate the correct insertion and the locus integrity. Non-specific integration of the donor DNA was characterized by two simple PCR amplifications using two sets of primers on the backbone of the plasmids.

To distinguish homozygous from heterozygous mice, tail DNA from F2 offspring was further characterized by PCR with three primers, two primers on the targeting genomic sequences and one primer on the transgene. Three independent *APOE3* or *APOE3S/S* lines without non-specific integration were selected and maintained. One *APOE3* or *APOE3S/S* line was used to cross with a transgenic mouse model overexpressing human tau harboring the P301S mutation (The Jackson Laboratory, 008169) to evaluate the effect of APOE3 Christchurch on tau pathology. Mice were housed no more than five per cage, given ad libitum access to food and water and housed in a pathogen-free barrier facility at 21–23 °C with 30–70% humidity on a 12-h light/12-h dark cycle. All mouse protocols were approved by the Institutional Animal Care and Use Committee at Weill Cornell Medicine. Specific primers for genotyping can be found in the STAR methods of this manuscript.

### Electrophysiological recording in freely moving mice

All experimental procedures were approved by the Weill Cornell Medical College Animal Care and Use Committee following National Institutes of Health guideline. Mice were initially anesthetized with 3.5 % isoflurane mounted on a stereotaxic frame and maintained under ∼1.2% isoflurane. Body temperature was maintained at 37 °C with a regulated heating blanket (World Precision Instruments, Sarasota, FL). A craniotomy was drilled for electrode insertion. Animals were implanted with a custom 16-ch 3D electrode array (Kedou Brain-Computer Technology, Suzhou, China) that contains 4 tetrodes over somatosensory (S1), the CA1 of hippocampus, dentate gyrus (DG), and primary visual cortex (V1) (S1 : antero-posterior (AP) -1.8mm, mediolateral (ML) 2.3mm , dorsoventral (DV) 1.0mm, CA1: AP -1.8mm, ML 1.5mm, DV1.4 mm, DG: AP -1.8mm, ML 1.0mm, DV 2.0mm, V1: AP -3.0mm, ML 2.3mm, DV 0.6mm. Fig xA). Stainless steel screws were implanted into the skull to provide electrical ground and mechanical stability for drives and the whole construct was bonded to the skull using C&B-Metabond luting cement (Parkell, Edgewood, NY). Two weeks after implantation, animals were put in a circular open field chamber (8228, Pinnacle Technology, Lawrence, KS) to record spontaneous activity continuously for 30 min. Electrophysiological data were acquired using an Intan RHD eadstage (Intan Technologies, LA, CA) and the Open Ephys acquisition board and software (OEPS, Alges, Portugal) sampled at 30kHz^64^. Locomotion was simultaneously acquired with a FLIR camera (Teledyne FLIR, North Billerica, MA) and the open-source Bonsia software^65^ recording at 50 Hz.

### Local field potential analysis

The offline LFP analysis was performed using custom-written Matlab script (MathWorks, Natick, MA). Briefly, LFP data will be preprocessed by first applying an anti-aliasing lowpass (<400Hz) zero-phase 8-order Chebyshev Type I filter then downsampling to 1000Hz. The power of theta (4-10 Hz) and gamma (30-90 Hz) were calculated then averaged by each tetrode to obtain the low and high band oscillation activity from each brain region. The normalized power was converted from the above value by dividing the average 10 sec baseline value at resting state when mice didn’t show any movement in the behavior video. The averaged power for the entire open field session was also calculated for the group comparison. Markless pose estimation for locomotion measurement was performed by deep learning based DeepLabCut from the behavioral video data set^66^.

### cFOS Mapping Experiment

8-9 month old male mice were habituated in single-house cages for 4 hours prior to start of experiment. Then, mice were individually placed in the aforementioned circular open field chamber, where their behavior was recorded from a camera underneath them. After 10 minutes of free-moving behavior recording, mice were placed back in their home cages for 45 minutes where they were promptly transcardially perfused and whole brains were collected for further 3D analysis.

### cGAS-inhibitor target engagement studies

To assess target engagement of TDI-6570 acutely, we administered TDI-6570 at a dose of 25mg/kg dissolved in DMSO via oral gavage to 8-9 month old mice. At 6 hours post-gavage, transcardial perfusion was performed and we collected tissue for both biochemical and immunohistochemical analyses.

### Tau seeding and spreading assay

To assess if the cGASi affects tau seeding and spreading, we injected 3.5-4.5 month old male and female *P301S* mice with 2uL of 1.8mg/ml K18 tau fibrils into their one hippocampi (anterior-posterior −2.0, medial-lateral +/− 1.3, dorsal-ventral −2.1). Mice were treated daily with cGASi (25 mg/kg body weight) or vehicle control mixed with almond butter Monday -Friday, and 150 mg/Kg chow diet over the weekend. Mouse weights and food consumed per c. age were collected throughout the study. After one month, mice were collected using transcardial perfusion (see methods below) and whole brains were collected for immunohistochemistry (see methods below) of MC-1 on the ipsilateral (seeding side) and contralateral (spreading side).

### Chronic cGAS inhibitor treatment

TDI-6570 were prepared using the methods as described previously^44^. Research diet pellets containing TDI-6570 (300 mg of drug per kg diet) were prepared by Research Diet (RDI) according to previously published methods, and control diet pellets were prepared similarly without TDI-6570^41^. The diet pellets were used before the expiration dates tentatively set for 6 months after the date of diet preparation. 6-7 month old male mice from all experimental conditions were treated with control or TDI-6570 pellets for 3 months before transcardial perfusion. Mouse weights and food consumed per cage were collected throughout the study.

### Tissue collection

Mice were euthanized using FatalPlus and perfused with cold PBS following cardiac puncture for blood collection. Blood was collected in EDTA-coated tubes and spun down to isolate plasma. After complete perfusion, left hemibrain was fixed in formalin for 24-48 hours followed by storage in 30% sucrose. Right hemibrain was microdissected for hippocampus and frontal cortex, which were frozen in dry ice and stored at -80°C until further processing.

### Immunohistochemistry

Fixed hemibrain was sectioned in 30 um-thick sections via microtome and stored in cryoprotectant. 3-5 hippocampal sections were picked per animal and following rinses with PBST, underwent antigen retrieval using Reveal Decloaker (Biocare Medical). Sections were incubated overnight at 4°C in primary antibodies: PSD-95 (Millipore, 1:400), IBA-1 (Abcam, 1:250), STING (Cell Signaling, 1:200), NeuN (Millipore, 1:200), MC-1, a conformational specific antibody for aggregated tau, (Gift from Peter Davies lab, 1:2500), GFAP (Abcam, 1:500), GM-130 (1:200), MBP (1:400). Following Alexafluor secondary antibody incubation at 1:500, sections were rinsed with PBST and mounted using Prolong Gold Anti-Fade reagent with DAPI. Slides were sealed with clear nail polish and kept at 4°C. Immunofluorescence was visualized using either LSM880 laser scanning confocal microscope and 3-4 images per animal were captured at 40X in CA1 and CA3 regions of hippocampus. Whole hippocampal images were taken at 10X with 11-13 planes and 2x2 images were stitched. After image capture, images were separated by color channel and underwent background subtraction and thresholding. Signal intensity was measured via ImageJ. Intensity of 1-2 control sections stained only with secondary antibody was averaged and subtracted from all sections. Signal intensity was normalized to area, averaged across sections for each animal, and plotted in Prism. Experimenters performing imaging and quantification were blinded.

### IMARIS 3D Reconstruction

For the 3D reconstruction of microglia, we used Z-stack images containing 25 stacks at 40 magnification in the CA1 region of the hippocampus on the Zeiss LSM 880 confocal. Raw czi files were used for further analysis during IMARIS software (version 9.31, Oxford Instruments). First, IMARIS was used to reconstruct the microglial surface using filaments. We then graphed the measurements: segment terminal points (for branch length) and soma volume. We used mixed model analysis to account for multiple cells in each image and multiple hippocampal sections/animal.

### Whole brain clearing and imaging for cFos Mapping

Fixed mouse brains from aged AD genetic cohorts were delipidated with a modified Adipo-Clear protocol^67,68^. Briefly, perfusion fixed brain samples were washed with B1n buffer (H2O/0.1% Triton X-100/0.3 M glycine, pH 7), then transferred to a methanol gradient series (20%, 40%, 60%, 80%) in B1n buffer, 4 mL for each brain, 1 h for each step; then 100% methanol for 1 h; then overnight incubation in 2:1 mixture of DCM:methanol and a 1.5 h incubation in 100% DCM the following day; then 100% methanol for 1 h three times, and reverse methanol gradient series (80%, 60%, 40%, 20%) in B1n buffer, 30 min for each step. Samples were then washed in B1n buffer for 1 h and overnight. The above procedures were done at room temperature with rocking to complete delipidation. The delipidated samples were then blocked in PTxwH buffer (PBS/0.1% Triton X100/0.05% Tween 20) with 5% DMSO and 0.3M glycine for 3 h and overnight at 37°C, then washed with PTxwH for 1 h, 2 h, and overnight at room temperature. For staining, brain samples were incubated in primary antibody (monoclonal rabbit-anti-cFOS, clone 9F6, Cell Signaling 2250S, Lot#11, 1:200) diluted in PTxwH for 2 weeks at 37°C. After primary antibody incubation, samples were washed in PTxwH for 1 h, 2 h, 4 h, overnight, then 1 d, and then incubated in secondary antibody (Jackson ImmunoResearch 711-607-003, Alexa Fluor 647 donkey-anti-rabbit, Lot#149379, 1:200) diluted in PTxwH at room temperature for 2 weeks. Samples were then washed in PTxwH for 1 h, 2 h, 4 h, overnight, then 1 d. Samples were further fixed in 1% PFA at 4°C overnight, washed in PTxwH at RT (1h, 2h, 4h, overnight), then blocked in B1n at RT overnight, and washed in PTxwH at RT (1h, 2h, 4h). Samples were then bleached in 0.3% H2O2 at 4°C overnight, washed 3 times in 20mM PB (16mM Na2HPO4, 4mM NaH2PO4 in H2O) at RT for 2 hours. For clearing, samples were dehydrated with methanol gradient with water (20%, 40%, 60%, 80%) 1h each step, then 100% methanol for 1h, DCM/methanol mixture overnight, and 100% DCM for 1h twice the next day, then 100% methanol for 1h twice and methanol gradient with water (80%, 60%, 40%, 20%) 1h each step. Next, brains were washed twice (1h, overnight) in 20mM PB buffer and finally twice in PTS solution (25% 2,2’-thiodiethanol/10mM PB) (3h, overnight), then equilibrated with 75% histodenz buffer (Cosmo Bio USA AXS-1002424) with refractive index adjusted to 1.53 using 2,2’-thiodiethanol. Samples were stored at -20°C until acquisition. The cleared brain samples were imaged horizontally with tiling using the LifeCanvas SmartSPIM lightsheet microscope. 647 nm lasers were used for the GFP viral labeling IHC imaging with the 3.6×/0.2 detection lens. Lightsheet illumination is focused with NA 0.2 lens, and axially scanned with electrically tunable lens coupled to the camera (Hamamatsu Orca Back-Thin Fusion) in slit mode. Camera exposure was set at fast mode (2 ms) with 16b imaging. The *X*/*Y* sampling rate was 1.866 μm and *Z* step set at 2 μm.

### Registration for Whole-brain imaging

Whole-brain 3D datasets were registered using the Common Coordinate Framework (CCF) reference brain, as previously described^69,70^. After downsampling full-resolution datasets, registration was performed in two steps: a 3D affine transformation followed by a 3D B-spline. Advanced Mattes Mutual Information was used to compute the similarity for both affine and B-spline transformations. The optimizer used affine transformation was Adaptive Stochastic Gradient Descent, and Standard Gradient Descent for B-spline transformation. After registration, the output transformations were used to transform the annotation atlas. Registration was performed using Elastix registration toolbox^71^.

### c-Fos cell segmentation and cell counting

The whole-brain signal distribution of c-Fos cells was automatically detected using custom Python scripts. First, the signal background was reduced using a rolling ball algorithm, and the signal threshold was set using adaptive thresholding. The centroids were calculated using 3D-connected components, and area-based counting was performed using the ARA atlas, as previously described. Statistical analysis was performed using ANOVA, with animal genotype as the independent variable and area counts as the dependent variables, followed by Tukey *post hoc* analysis.

### Single nuclei RNA sequencing

Nuclei isolation from frozen mouse hippocampi was adapted from a previous study [Habib 2017], with modifications. All procedures were performed on ice or at 4°C. In brief, postmortem brain tissue was placed in 1,500 µl of Sigma nuclei PURE lysis buffer (Sigma, NUC201-1KT) and homogenized with a Dounce tissue grinder (Sigma, D8938-1SET) with 15 strokes with pestle A and 15 strokes with pestle B. The homogenized tissue was filtered through a 35-µm cell strainer, centrifuged at 600g for 5 min at 4 °C and washed three times with 1 ml of PBS containing 1% bovine serum albumin (BSA, Thermo Fisher Scientific, 37525), 20 mM DTT (Thermo Fisher Scientific, 426380500) and 0.2 U µl^−1^ recombinant RNase inhibitor (Ambion, AM2684). Nuclei were then centrifuged at 600g for 5 min at 4 °C and resuspended in 350 µl of PBS containing 0.04% BSA and 1× DAPI, followed by fluorescence-activated cell sorting to remove cell debris. The sorted suspension of DAPI-stained nuclei was counted and diluted to a concentration of 1,000 nuclei per µl in PBS containing 0.04% BSA.

For droplet-based snRNA-seq, libraries were prepared with Chromium Single Cell 3′ Reagent Kits v3 (10x Genomics, PN-1000075) according to the manufacturer’s protocol. The snRNA-seq libraries were sequenced on a NovaSeq 6000 sequencer (Illumina) with 100 cycles. Gene counts were obtained by aligning reads to the mm10 genome with Cell Ranger software (v.3.1.0; 10x Genomics). To account for unspliced nuclear transcripts, reads mapping to pre-mRNA were counted. Cell Ranger 3.1.0 default parameters were used to call cell barcodes. We further removed genes expressed in no more than three cells, cells with unique gene counts over 4,000 or less than 300, cells with UMI counts over 20,000 and cells with a high fraction of mitochondrial reads (>5%). Potential doublet cells were predicted using DoubletFinder for each sample separately, with high-confidence doublets removed. Normalization and clustering were done with the Seurat package v3.2.2 (Stuart, Butler 2019). In brief, counts for all nuclei were scaled by the total library size multiplied by a scale factor (10,000) and transformed to log space. A set of 2,000 highly variable genes was identified with SCTransform from the sctransform R packageFindVariableFeatures function with vst method. This returned a corrected unique molecular identifier count matrix, a log-transformed data matrix and Pearson residuals from the regularized negative binomial regression model. Principal-component analysis was done on all genes, and t-distributed stochastic neighbor embedding was run on the top 15 principal components. Cell clusters were identified with the Seurat functions FindNeighbors (using the top 15 principal components) and FindClusters (resolution = 0.1). In this analysis, the neighborhood size parameter pK was estimated using the mean variance-normalized bimodality coefficient (BCmvn) approach, with 15 principal components used and pN set as 0.25 by default. Sample integration was performed using FindIntegrationAnchors and IntegrateData functions in Seurat. For the baseline *E3, E3P301S, E3^S/S^ and E3^S/S^P301S* cohort snRNAseq, we sequenced 3 hippocampi/group. For the cGASi-treated cohort, we also sequenced 3 hippocampi/group for the following groups: *E3/control diet, E3^S/S^/control diet, E3P301S/control diet, E3^S/S^P301S/control diet, E3/cGASi, E3^S/S^/cGASi, E3P301S/cGASi.* For each cluster, we assigned a cell-type label using statistical enrichment for sets of marker genes and manual evaluation of gene expression for small sets of known marker genes. Differential gene expression analysis was done using the FindMarkers function and MAST (Finak 2015). To identify gene ontology and pathways enriched in the DEGs, DEGS were analyzed using the MSigDB gene annotation database (Subramanian 2005, Liberzon 2011). To control for multiple testing, we used the Benjamini–Hochberg approach to constrain the FDR.

### Western blot

25 ug frontal cortex or hippocampal lysate prepared above were boiled for 5 minutes and run on 26-well 4-12% Bis-Tris gels (Invitrogen) using MES buffer (Invitrogen) for 1 hour. Proteins were transferred from gel onto PVDF membrane for 2 hours. Membranes were washed three times for 10 min each in TBS with 0.01% triton X-100 (TBST). Membranes were blocked for 30 minutes in 5% milk in TBST and incubated with APOE (1:800, CalBiochem), STING (1:1000), pTBK (1:1000), TBK (1:1000) and GAPDH (1:1,000, GeneTex) antibodies overnight in cold room. The following day, membranes were washed three times for 10 min each in TBST and incubated in appropriate HRP secondary for 1 hour, rinsed, and followed by ECL development and imaging using Bio-Rad imager.

### Lipidomic analyses: Lipid extraction

For lipidomic analysis, hippocampal samples from 6 mixed sex mice were homogenized in with ice-cold milli-Q water containing a cOmplete, Mini Protease Inhibitor Cocktail using a bead mill homogenizer (VWR). Protein concentration in the lysates was quantified using the BCA assay, and approximately 50 μg of tissue lysate was transferred to Pyrex glass tubes with PTFE-lined caps. Lipid extraction followed the Folch method^72^: 6 mL of ice-cold chloroform and methanol (2:1 v/v) and 1.5 mL of water were added to each sample. The tubes were thoroughly vortexed to ensure homogeneous mixing of polar and non-polar solvents. SPLAH internal standards were added before extraction. The samples were centrifuged at 1000 rpm for 20 minutes at 4°C to separate the organic and aqueous phases. The lower organic phase was carefully pipetted into a new glass tube using a sterile glass pipette, avoiding the intermediate layer containing cell debris and precipitated proteins. The organic phase was then dried under a nitrogen stream until all solvents evaporated. Finally, the samples were reconstituted in 150 μL of chloroform:methanol (2:1) and stored at -80°C until mass spectrometry (MS) analysis.

### Gangliosides extraction

The aqueous layer containing gangliosides was collected and dried under a gentle stream of nitrogen. The dried aqueous phase was reconstituted in 1 mL pure water and subsequently desalted by using Sola HRP SPE 30mg/2mL 96-well plate 1EA (Thermo Scientific #60509-001). Initially, the cartridges were cleaned 3 times with 1 mL of MeOH and equilibrated 3 times with water. Samples were loaded onto the column, washed 3 times with water and, finally, the gangliosides were eluted by three times 1 mL of MeOH. The eluate was dried under nitrogen flow and reconstituted in MeOH/H2O/CHCl3 (60:9:120, v/v/v).

### Lipidomic analyses: LC-MS/MS analyssis of lipidomics

Lipids were separated using ultra-high-performance liquid chromatography (UHPLC) coupled with tandem mass spectrometry (MS/MS). UHPLC analysis was conducted on a C30 reverse-phase column (Thermo Acclaim C30, 2.1 x 150 mm, 2.6 μm) maintained at 50°C and connected to a Vanquish Horizon UHPLC system, along with an OE240 Exactive Orbitrap MS (Thermo Fisher Scientific) equipped with a heated electrospray ionization probe. Each sample (2 μL) was analyzed in both positive and negative ionization modes. The mobile phase included 60:40 water:acetonitrile with 10 mM ammonium formate and 0.1% formic acid, while mobile phase B consisted of 90:10 isopropanol:acetonitrile with the same additives. The chromatographic gradient involved: Initial isocratic elution at 30% B from -3 to 0 minutes, followed by a linear increase to 43% B (0-2 min), then 55% B (2-2.1 min), 65% B (2.1-12 min), 85% B (12-18 min), and 100% B (18-20 min). Holding at 100% B from 20-25 min, a linear decrease to 30% B by 25.1 min, and holding from 25.1-28 min. Flow rate of 0.26 mL/min, injection volume of 2 μL, and column temperature of 55°C. Mass spectrometer settings included an ion transfer tube temperature of 300°C, vaporizer temperature of 275°C, Orbitrap resolution of 120,000 for MS1 and 30,000 for MS2, RF lens at 70%, with a maximum injection time of 50 ms for MS1 and 54 ms for MS2. Positive and negative ion voltages were set at 3250 V and 2500 V, respectively. Gas flow rates included auxiliary gas at 10 units, sheath gas at 40 units, and sweep gas at 1 unit. High-energy collision dissociation (HCD) fragmentation was stepped at 15%, 25%, and 35%, and data-dependent tandem MS (ddMS2) ran with a cycle time of 1.5 s, an isolation window of 1 m/z, an intensity threshold of 1.0e4, and a dynamic exclusion time of 2.5 s.

Full-scan mode with ddMS^2^ was performed over an m/z range of 250-1700, with EASYICTM used for internal calibration. The raw data were processed and aligned with LipidSearch 5.1, using a precursor tolerance of 5 ppm and a product tolerance of 8 ppm. Further filtering and normalization were conducted using an in-house app, Lipidcruncher. Semi-targeted quantification was performed by normalizing the area under the curve (AUC) to the AUC of internal standards and further normalized with the total quantified protein level.

### For LC-MS/MS analysis of gangliosides

Samples were analyzed using a Vanquish UHPLC system (Thermo Scientific) coupled to an Orbitrap Exploris 240 mass spectrometer (Thermo Scientific, #BRE725535). Separation was achieved on a Kinetex HILIC column (Phenomenex, #00D-4461-AN; 2.6 μm, 100 x 2.1 mm).

The mobile phase consisted of solvent A (acetonitrile with 0.2% v/v acetic acid) and solvent B (water containing 10 mM ammonium acetate, pH 6.1, adjusted with acetic acid). The column temperature was maintained at 50 °C. A gradient elution was employed at a constant flow rate of 0.6 mL/min: 12.3% B at 0 min, a linear increase to 22.1% B from 1 to 15 minutes, followed by column equilibration at 12.3% B for 5 minutes.

Mass spectrometry analysis was performed in heated electrospray ionization (HESI) mode under the following conditions: spray voltage at −4.5 kV, heated capillary temperature at 300 °C, and vaporizer temperature at 250 °C. The gas settings were as follows: sheath gas at 40 units, auxiliary gas at 5 units, and sweep gas at 1 unit. The ion transfer tube temperature was maintained at 300 °C. For MS1 analysis, the Orbitrap resolution was set to 120,000 with a scan range of 700–1800 m/z, RF lens at 60%, and an AGC target set to standard. For MS2, the Orbitrap resolution was set to 30,000. Internal calibration was achieved using EASY-IC™. LipidSearch 5.1, using a precursor tolerance of 5 ppm and a product tolerance of 8 ppm. Further filtering and normalization were conducted using an in-house app, Lipidcruncher. Quantification was achieved by normalizing the area under the curve to GM3-d5 standards, followed by further normalization to the amount of protein used in the preparation.

### Primary microglial culture

Primary microglial cells were collected from mouse pups at postnatal days 1–3. Briefly, the brain cortices were isolated and minced. Tissues were dissociated in 0.25% Trypsin-EDTA for 10 min at 37 °C and agitated every 5 min. Two hundred microliters of DNAse I (Millipore) was then added. Trypsin was neutralized with complete medium (DMEM; Thermo Fisher) supplemented with 10% heat-inactivated FBS (Hyclone), and tissues were filtered through 70-μm cell strainers (BD Falcon) and pelleted by centrifugation at 250*g*. Mixed glial cultures were maintained in growth medium at 37 °C and 5% CO2 for 7–10 d *in vitro*. Once bright, round cells began to appear in the mixed glial cultures, recombinant mouse granulocyte– macrophage colony-stimulating factor (1 ng ml^−1^; Life Technologies) was added to promote microglia proliferation. Primary microglial cells were collected by mechanical agitation after 48–72 h and plated on poly-D-lysine-coated 24-well plates (Corning) in growth medium. Microglia were maintained in DMEM supplemented with 10% FBS, 100 U ml^−1^ penicillin and 100 μg ml^−1^ streptomycin. At 24 hours after plating, microglia were treated with 1ug/ml of ON4R recombinant tau obtained from Dr. Sue-Ann Mok’s group or Aβ monomers at a concentration of 4.2ug/mL from Dr. Yueming Li’s lab for 24 hours.

### Bulk sequencing analysis

Primary microglia isolated from *E3* and *E3^S/S^* pups were isolated and treated with tau or amyloid as described above and collected 24 hours later. Total RNA was extracted from the samples using QuickRNA MicroPrep Kit (Zymo Research). After RNA isolation, total RNA integrity was checked using a 2100 Bioanalyzer (Agilent Technologies), and concentrations were measured by Nanodrop (Thermo Fisher. RNA was isolated from microglia using the Quick-RNA MicroPrep Kit (Zymo Research, R1051). RNA was shipped to Novogene for library preparation and bulk RNA sequencing. Differential gene expression was analyzed with the DESeq2 1.38.3 package131. Counts were normalized using the median of ratios method. Genes with <15 counts across all samples were excluded from analysis. To control for multiple testing, we used the Benjamini–Hochberg approach to constrain the FDR. Pathway analysis was done using the MSigDB gene annotation database. Ingenuity Pathway Analysis (QIAGEN, Inc.) was used to identify gene activation networks and upstream regulators. Raw read counts per gene were extracted using HTSeq-count v0.11.2^73^.

### Tau pulse-chase experiment

For pulse-chase experiments, primary microglia were plated after shaking onto 8-well chamber slides at a density of 1 x 10^5^ cells/well. 24 hours after plating, media with 1ug/ml of 0N4R tau fibrils from Dr. Sue-Ann Mok’s lab was added to each well or fresh media was changed for control wells. After 2 hours, media was collected from both conditions and ½ of the wells were washed 3 times with PBS and fixed for 15 minutes with 4% PFA and sucrose. After 15 minutes, cells were washed 3 times with PBS. The remaining wells were exchanged with fresh media for another 22 hours. At 24 hours post-initial tau treatment, media was collected and remaining cells were fixed as previously described. Fixed chamber wells were stored at 4C until immunocytochemistry for tau was performed. Cells were washed with PBS and blocked for 1 hour at room temperature with 10% BSA in TBST. After washing with PBS again, cells were incubated with HT-7 tau antibody (1:600) diluted in 1% BSA in TBST) for 2 hours at room temperature, then washed 3 times, and incubated with Alexa 568 secondary antibody (1:500) for 1 hour at room temperature. Coverslips were placed with anti-fade reagent containing DAPI and sealed with nail polish.

### Human iPSC-derived microglia-like cell culture and differentiation

Human *APOE3* and its isogenic *APOE3 Christchurch* iPSCs were obtained by Jackson Laboratory. These iPSCs were cultured with mTeSR plus media (STEMCELL) in hESC-Qualified Matrigel (Corning)-coated 6-well plates. At 70∼80% confluence, the iPSCs were dissociated with ReLeSR (STEMCELL) solution. The dissociated iPSCs were seeded in to new Matrigell-coated-6-Well plate with mTeSR plus media containing 10 μM ROCK inhibitor (Tocris). For generating the human HPCs, we used the STEMdiff Hematopoietic kit (STEMCELL). Briefly, dissociated iPSCs with ReLeSR were seeded in Matrigel-coated-6-well plate with mTeSR plus containing 10 μM ROCK inhibitor. At Day 0, mTeSR plus media was replaced with Basal media containing Sup A. At Day 3, the media was replaced with Basal media containing Sup B. Half of the media was changed with Sup B containing media every other day. At Day 13, we used HPCs for Microglia like-cell differentiation. To generate iMGLs, we used Microglia Basal media [DMEM/F12 (Gibco), B27 (Gibco), N2 (Gibco), GlutaMAX (Gibco), NEAA (Sigma Aldrich), Monothioglycerol (Sigma Aldrich), insulin-transferrin-selenite (Gibco), and insulin (Sigma Aldrich)] 50,000 of HPCs were seeded in Matrigel-coated 6-well plate with Microglia Basal media containing 100 ng/ml of IL-34 (Peprotech), 50 ng/ml of TGF-β1 (STEMCELL), and 25 ng/ml or M-CSF (R&D Systems). 1ml of the Basal media containing 3 cytokines was added every other day until Day 24. On Day 25, the media was replaced with the media additionally containing 2 more cytokines [100 ng/ml of CD200 (Novo protein) and 100ng/ml of Cx3CL1 (STEMCELL)]. At Day 28, iMGLs were used for experiments.

### iMGL Tau Uptake Assay

iMGLs were seeded to a Poly-D lysine-coated 6-well plate at a density of 1.5 x 10^5^/well. After 48 hours of seeding the cells, 1 μg/ml of 0N4R tau fibrils from Dr. Sue-Ann Mok’s lab was treated to iMGLs. After 24 hours, the level of tau in the media was measured using a human total tau ELISA kit (Invitrogen). The level of tau was normalized with protein concentration. How about the levels of intracellular tau?

### IP-10 ELISA

To measure the secreted IP-10 from iMGLs after tau treatment, we used human IP-10 ELISA kit (Invitrogen). iMGLs were seeded in a Poly-D lysine-coated 6-well plate at a density of 1.5 x 10^5^/well. After 48 hours, 1 μg/ml of tau was treated to iMGLs. After 24 hours of incubation, the human IP-10 from iMGLs was measured using a human IP-10 kit. The level of IP-10 was normalized with protein level.

## SUPPLEMENTARY MATERIALS

Figs. S1–S7

Tables S1 to S5

**Figure S1.**
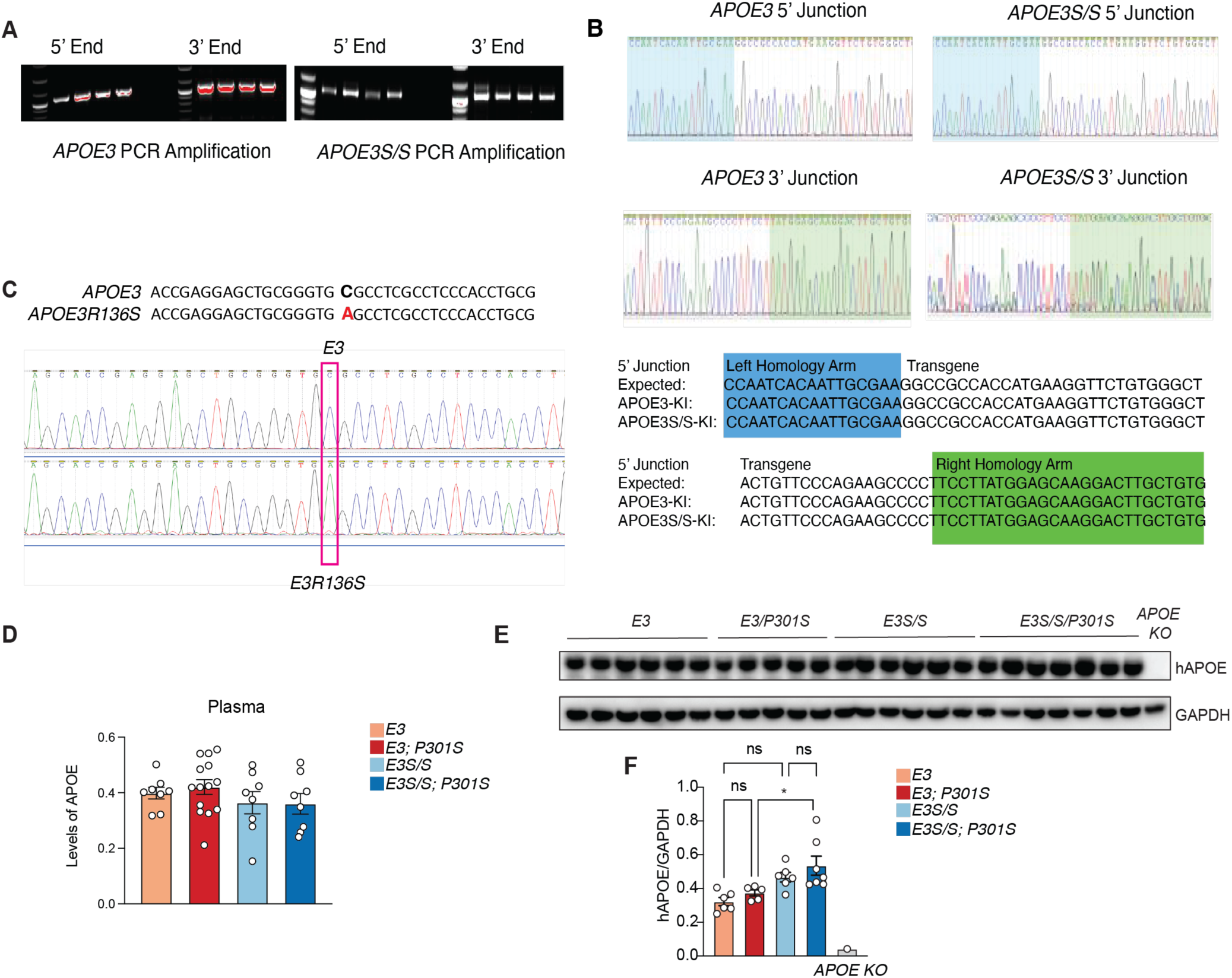
Sequencing confirmation and APOE expression levels of *ApoE3* and *ApoE3^S/S^* knock-in mice. Related to Figure 1. (A) Representative gel for PCR amplification to show insertion of the *E3* and *E3^S/S^*. (B) Sequence of human APOE3-KI and human APOE3S/S-KI. (C) Confirmation of *R136S* mutation using Sanger sequencing. (D) Plasma levels of APOE collected from all four experimental groups at 9-10 months in male mice, also showing no significant differences between experimental groups. (E) Representative western blot of human APOE levels in frontal cortex lysates of female mice from each experimental group, showing no differences between any group. (F) Quantification of APOE levels in frontal cortex lysates of female mice using APOE KO as control, normalized to GAPDH, showing no differences among any experimental groups. Data are reported as mean ± SEM. *p<0.05 for *E3/P301S* compared to *E3^S/S^/P301S*. Data were analyzed by one-way ANOVA with Tukey’s multiple comparison test.

**Figure S2.**
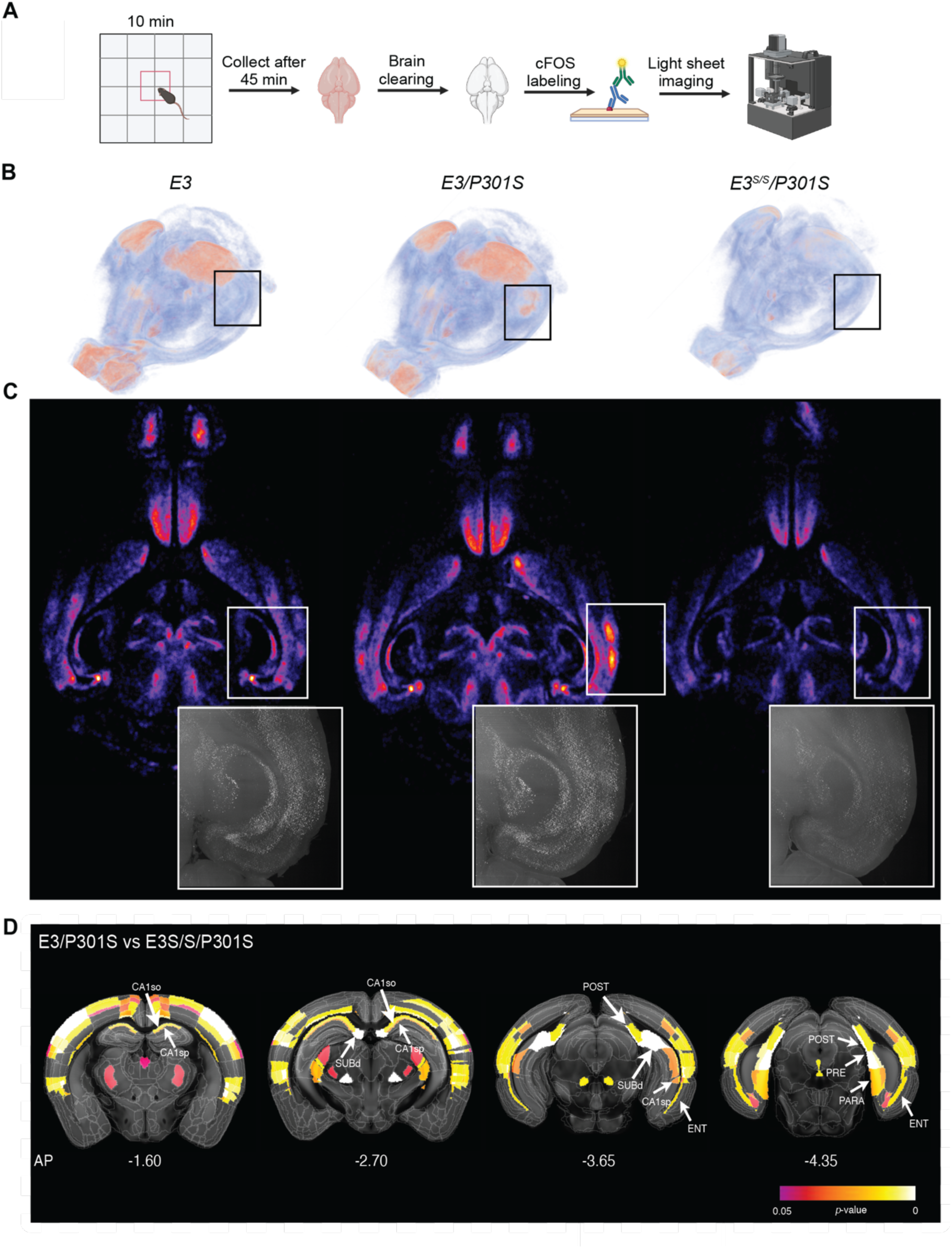
3D c-Fos mapping reveals *E3^S/S^* protects against tau-induced circuit alterations in hippocampus and entorhinal cortex. Related to Figure 2. (A) Schematic of experimental design. (B) 3D perspective view (OB pointing bottom-left) of voxelated whole-brain c-Fos activity in 9-10 month old male E3, *E3P301S* and *E3^S/S^P301S* mice. Boxed region indicates the HPC-EC area. (C) Middle horizontal plane (OB pointing up) of voxel-wised c-Fos activity in E3, *E3P301S,* and *E3^S/S^P301S* brains. The dashed line points to the hippocampal formation, showing tau-induced c-Fos hyperactivity in *E3P301S* and suppression in *E3^S/S^P301S*. The lower zoom-in panel shows a 500 mm optical maximum-intensity projection view of the raw c-Fos labeling pattern in the same hippocampal formation area. (D) Coronal atlas planes (AP axis: -1.60, -2.70, -3.65 and -4.35) covering major hippocampal formation area, with color-coding showing the brain anatomical regions with increased c-Fos induced activity in *E3P301S* compared with *E3^S/S^P301S*. Significant brain anatomical regions are color-coded based on the statistical *p-*value. Arrows point to the main hippocampal formation regions with increased activity in *E3/P301S* compared with *E3^S/S^P301S*: CA1, pyramidal layer (CA1sp); CA1, stratum oriens (CA1so); subiculum, dorsal part (SUBd); postsubiculum (POST); parasubiculum (PARA); presubiculum (PRE); entorhinal cortex (ENT).

**Figure S3.**
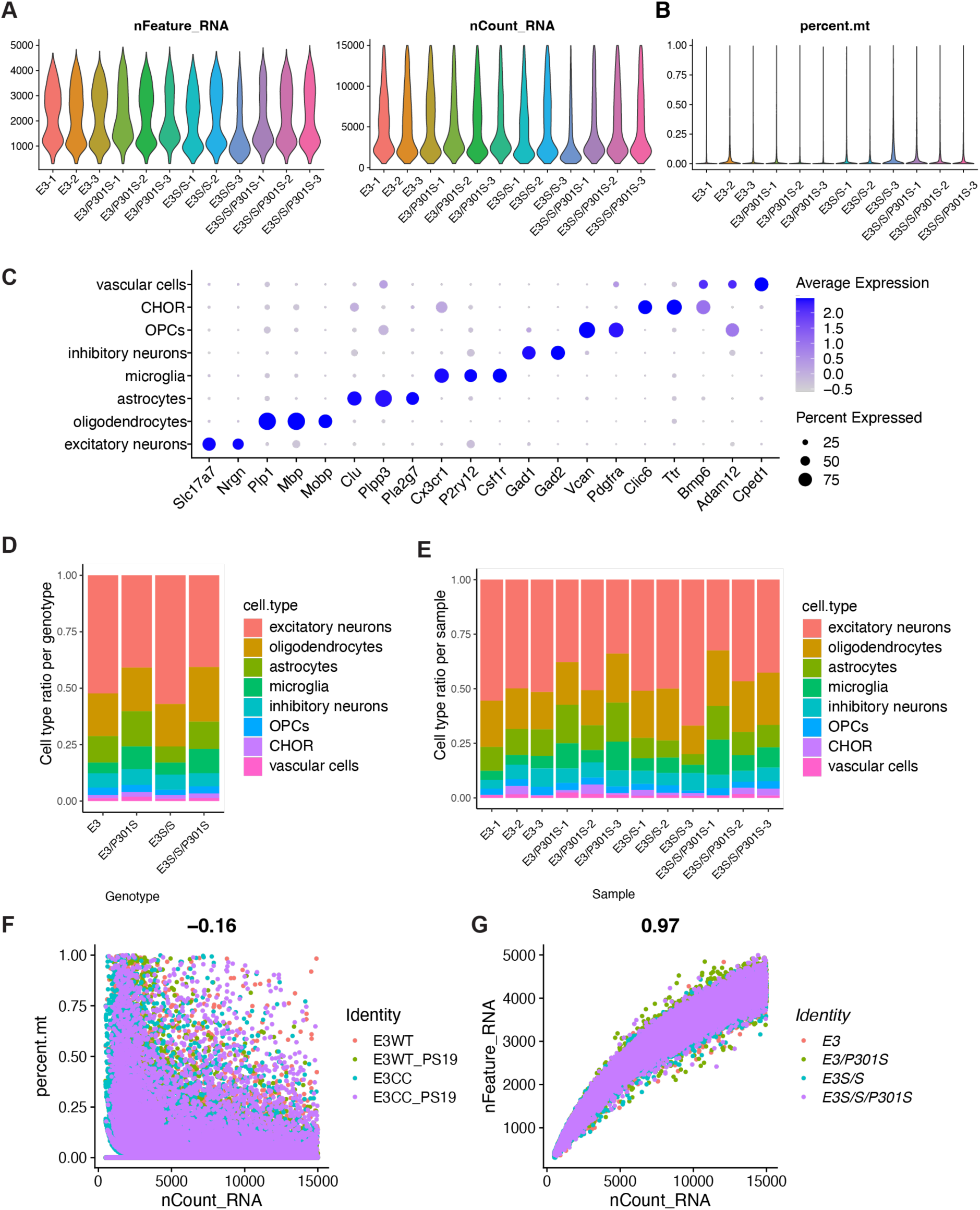
Quality control assessment of single nuclei RNA sequencing data of the E3 and E3S/S tauopathy mice, related to Figures 3, 4, 5. (A) Quality-control plots showing equivalent amounts of total RNA features, total number of RNA counts. (B) Violin plot showing percentage of mitochondrial genes detected per nuclei for each individual sample. (C) Dot plot showing expression of identity markers for each cell type. (D) Proportion of each cell type within each genotype. (E) Proportion of each cell type within individual samples. (F and G) Correlation between UMI counts and percentage of mitochondrial genes detected (F) or total gene counts (G) per nuclei for each individual sample.

**Figure S4.**
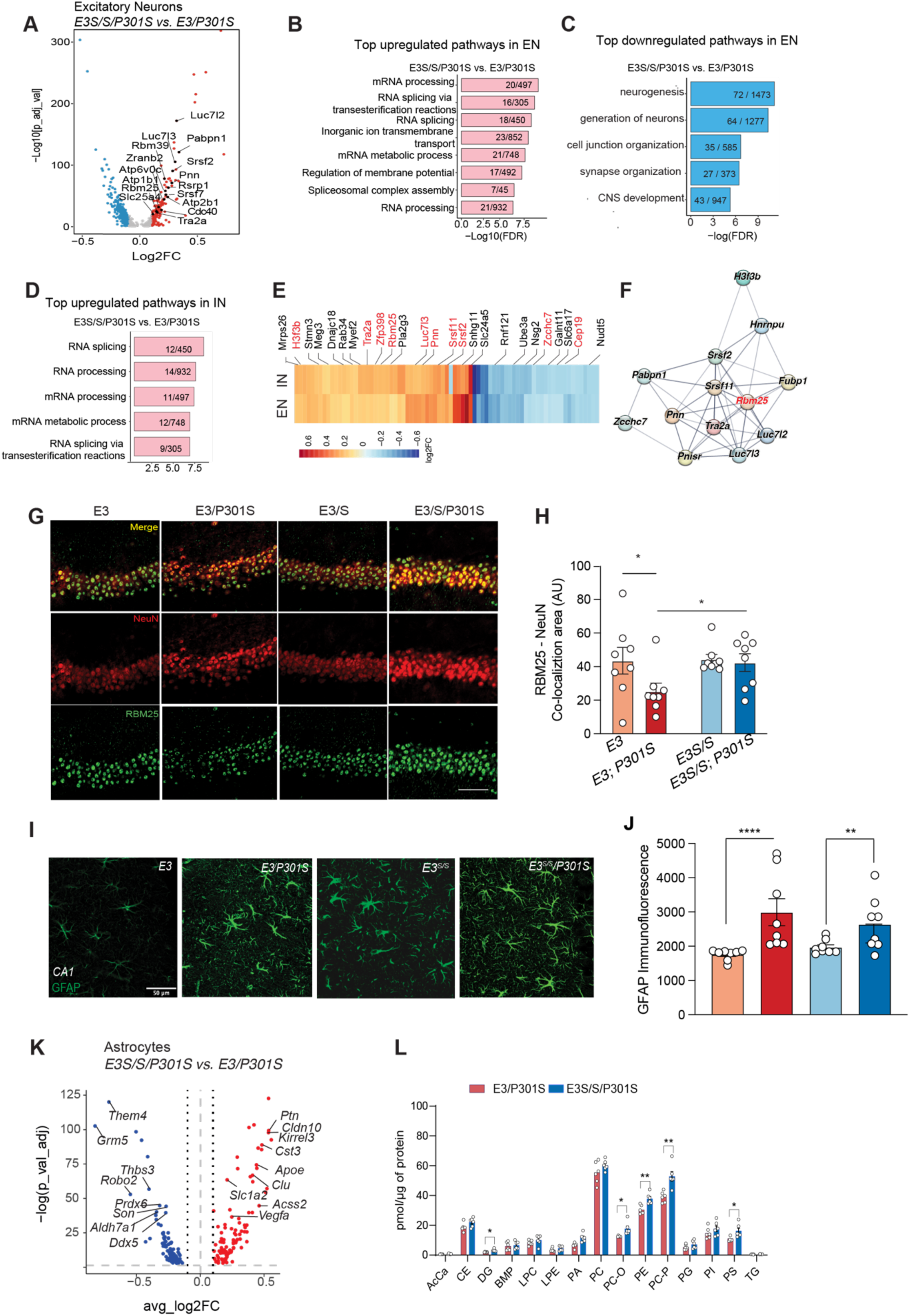
R136S mutation alters astroglial and neuronal transcriptomes in hippocampus of tauopathy mice. Related to Figure 3. (A) Volcano plot of DEGs in excitatory neurons between *E3^S/S^/P301S* and *E3/P301S.* Dashed lines represent log2foldchange threshold of 0.1 or -0.1and adjusted p value threshold of 0.05. (B) Top upregulated pathways in excitatory neurons between *E3^S/S^/P301S* and *E3/P301S* using GSEA. (C) Top downregulated pathways in excitatory neurons between *E3^S/S^/P301S* and *E3/P301S* using GSEA. (D) Top upregulated pathways in inhibitory neurons between *E3^S/S^/P301S* and *E3/P301S* using GSEA. (E) Heatmap of top 50 upregulated and downregulated genes of excitatory and inhibitory neurons between *E3^S/S^/P301S* and *E3/P301S,* showing splicing related genes in red. (F) String analysis showing RNA splicing related genes in neurons. (G) Representative immunofluorescence images of RBM25 (green) and neuronal marker, NeuN (red) and merged (yellow) in all four experimental groups. (H) Quantification of colocalization of RBM25 and NEUN signal across genotypes, n=8 mice per genotype with 2-3 sections per mouse. Data are reported as mean ± SEM and analyzed via two-way ANOVA with mixed effects model. *p<0.05. (I) Representative immunofluorescence images of GFAP (green) in CA1 subregion of hippocampus in mice across four genotypes; scale bar, 50um. (J) Quantification of GFAP immunofluorescence intensities, showing increased GFAP in both *P301S* groups. Results are presented as average intensity measures from 3-4 sections per animal with 8 animals/experimental group. Data are reported as mean ± SEM. ****p<0.0001 for *E3* compared to *E3/P301S* and **p=0.0073 for *E3^S/S^* compared to *E3^S/S^/P301S*. Data were analyzed by two-way ANOVA with mixed-effects model. (K) Volcano plot of DEGs in astrocytes between *E3^S/S^/P301S* and *E3/P301S*. Dashed lines represent log2foldchange threshold of 0.1 or -0.1 and adjusted p value threshold of 0.05. (L) The lipid composition in the E3S/S/P301S mouse brain is altered compared to the E3/P301S mice. Notable lipid changes include significantly elevated levels of PC-O, PE, and PE-P in the E3S/S/P301S group relative to E3/P301S. The lipid classes are presented as the mean of all lipid species measured, with error bars indicating the mean ± SEM. cCa: acylcarnitine, BMP: bis(monoacylglycerol)phosphate, CE: cholesterol ester, DAG: diacylglycerol, LPC: lysophosphatidylcholine, LPE: lysophosphatidylethanolamine, LPI: lysophosphatidylinositol, LPS: lysophosphatidylserine, PA: phosphatidic acid, PC: phosphatidylcholine, PC-O: alkyl-phosphatidylcholine, PE: phosphatidylethanolamine, PE(O): alkyl-phosphatidylethanolamines, PG: phosphatidylglycerol, PI: phosphatidylinositol, PS: phosphatidylserine, TAG: triacylglycerol.

**Figure S5.**
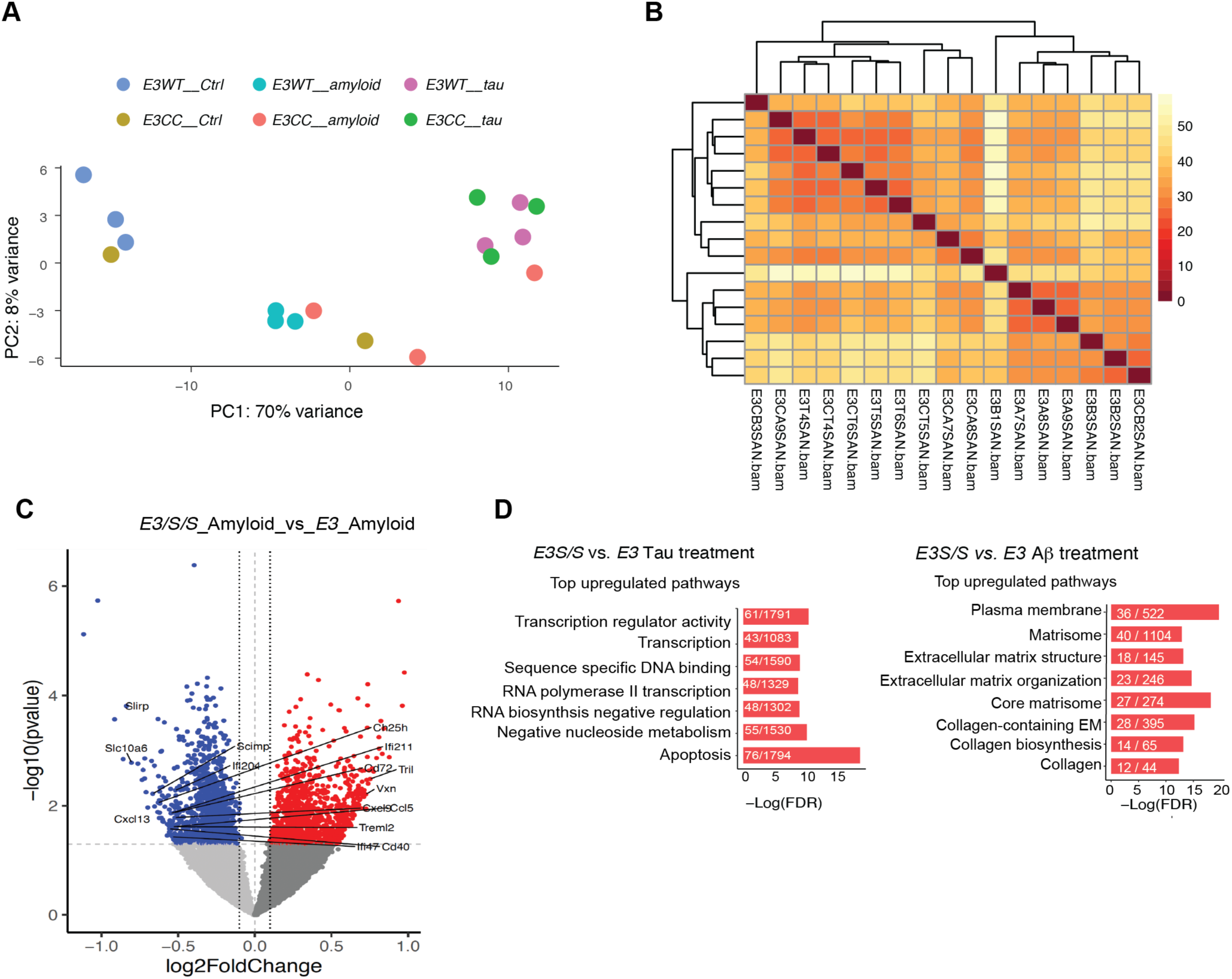
QC for primary microglia experiments and upregulated pathways by R136S mutation in the presence of tau and Aß, related to Figure 5. (A) Principal component analysis plot. Tau fibril stimulation accounts for 83% of gene expression variance. Each circle represents one biological sample. (B) Heatmap showing correlations between each biological sample. Samples cluster based on *APOE3* genotype and tau stimulation status. There are no outlier samples. (C) Volcano plot of DEGs in primary microglia following Aß stimulation between *E3^S/S^* and *E3*. Dashed lines represent log2foldchange threshold of 0.1 and adjusted p value threshold of 0.05. (D) Top upregulated pathways in tau-stimulated or amyloid-stimulated, respectively, *E3^S/S^* versus *E3* primary microglia using GSEA.

**Figure S6.**
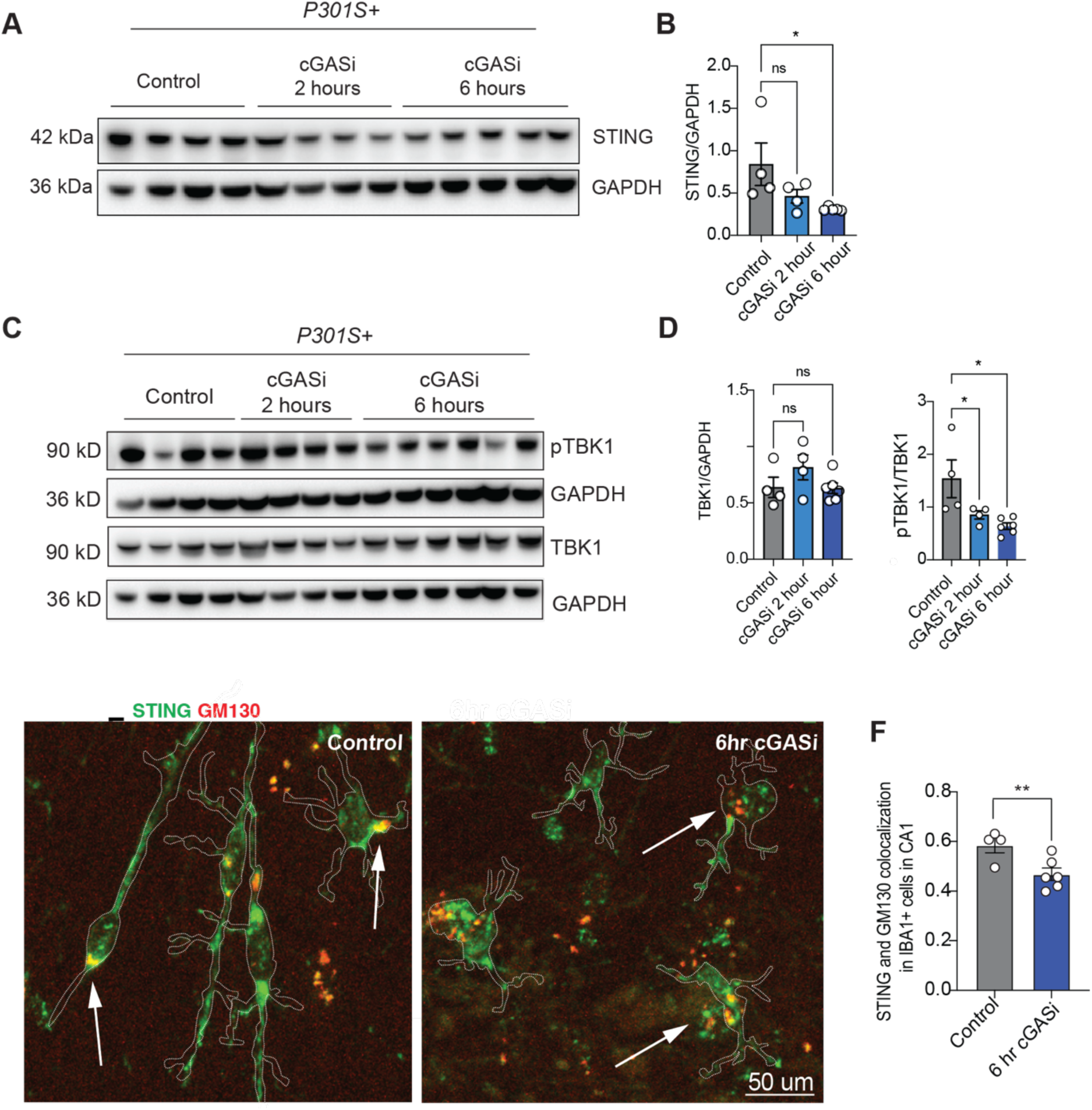
cGAS inhibitor enhances phagocytosis of tau in primary microglia and reduces STING and pTBK1 in tauopathy mice, related to Figure 7. (A) Western blot for STING and GAPDH of frontal cortex lysate of *P301S* mice treated with control, cGASi for 2 hours or 6 hours via oral gavage before collection. (B) Quantification of ratio of STING/GAPDH in frontal cortex lysates of control, cGASi 2 hour treatment and cGASi 6 hour treatment, showing significant reduction of STING/GAPDH at 6 hours post-cGASi treatment. Data are reported as mean ± SEM. *p<0.05 for control compared to 6hr cGASi treated mice . Data were analyzed by one-way ANOVA with Dunnett’s test. (C) Representative western blot for pTBK1, GAPDH and TBK1 of frontal cortex lysate of *P301S* mice treated with control, cGASi for 2 hours or 6 hours before collection. (D) Quantification of ratio of pTBK1/TBK1 in frontal cortex lysates of control, cGASi 2 hour treatment and cGASi 6 hour treatment, showing significant reduction of pTBK1/TBK1 at 2 hours post-cGASi treatment and more significant by 6 hours. Data are reported as mean ± SEM. Data were analyzed by one-way ANOVA with Dunnett’s test. (E) Representative fluorescence images of STING (green) and GM130 (red) colocalization in Iba1+ cells (outlined in white) in mice treated with control or cGAS inhibitor for 6 hours; scale bar = 25um. (F) Quantification of STING and GM130 colocalization in Iba1+ cells in control and cGASi treated mice, showing decreased colocalization after cGASi treatment. Results are presented as average intensity measures from 3-4 sections per animal with 4-6 animals/experimental group. Data are reported as mean ± SEM. **p=0.0029 for control treatment compared to cGASi treatment. Data were analyzed by two-way ANOVA with mixed-effects model.

**Figure S7.**
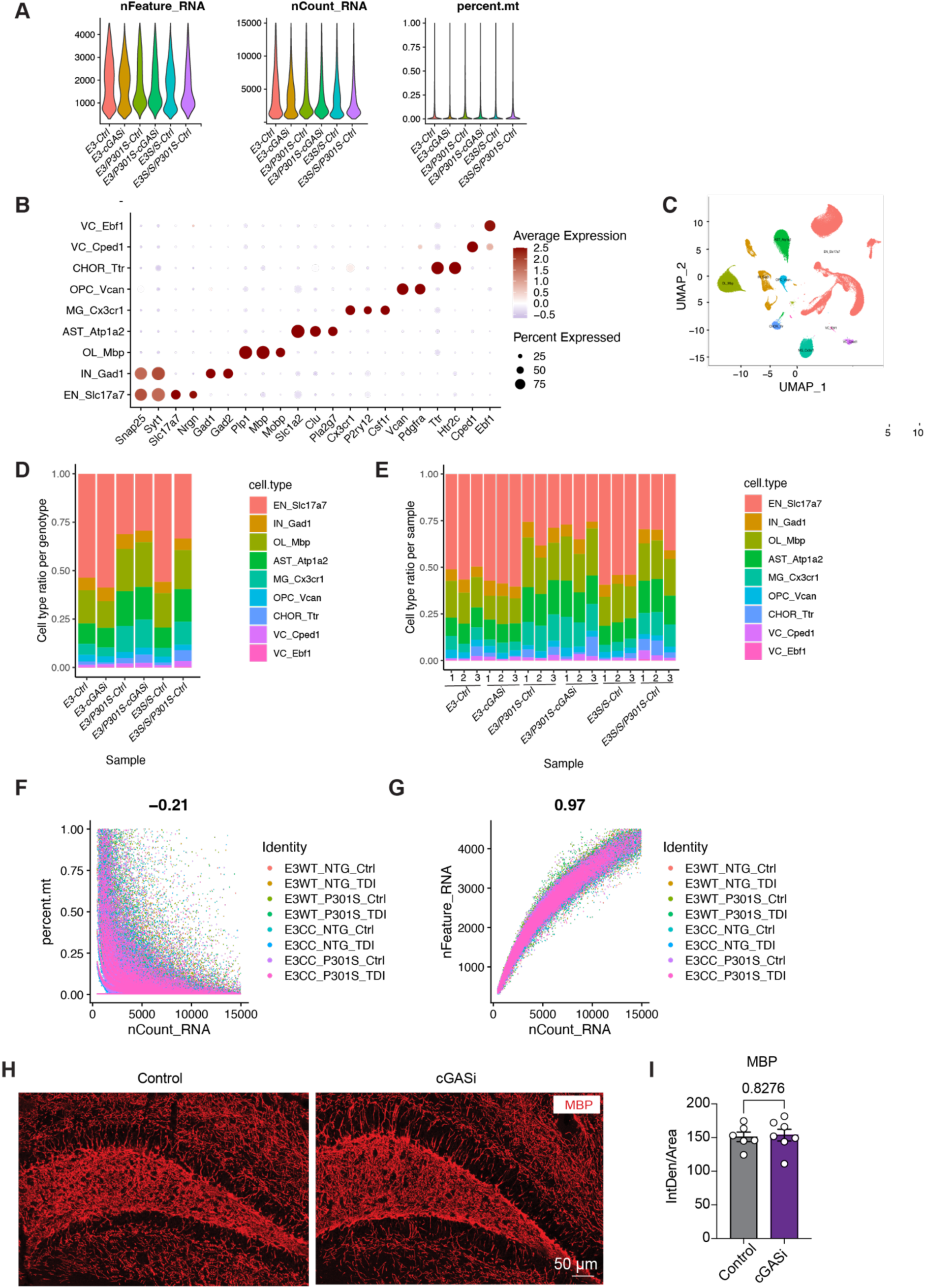
Quality control assessment of single nuclei RNA sequencing data of the *E3* and *E3^S/S^* tauopathy mice treated with cGASi and the effects of, related to Figure 7. (A) Quality-control plots showing equivalent amounts of total RNA features, total number of RNA counts, showing percentage of mitochondrial genes detected per nuclei for each individual sample. (B) Dot plot showing expression of identity markers for each cell type. (C) UMAP all cell types of all samples. (D) Proportion of each cell type within differing *E3* and *E3^S/S^* genotypes, treated with either cGAS-inhibitor diet or control diet. (E) Proportion of each cell type within individual samples. (F–G) Correlation between UMI counts and percentage of mitochondrial genes detected (E) or total gene counts (F) per nuclei for each individual sample. (H) Immunostaining for MBP in dentate gyrus of 9-10 month old male *P301S* mice treated with control diet or cGASi diet for 3 months in chow before collection. (I) Quantification of ratio of MBP signal in dentate gyrus of 9-10 month old male *P301S* mice treated with control diet or cGASi diet for 3 months in chow before collection, showing no difference in MBP levels between treatment groups. Data are reported as mean ± SEM. Data were analyzed by unpaired t test (two-tailed).

## REFERENCES

1. Arboleda-Velasquez, J.F., Lopera, F., O’Hare, M., Delgado-Tirado, S., Marino, C., Chmielewska, N., Saez-Torres, K.L., Amarnani, D., Schultz, A.P., Sperling, R.A., et al. (2019). Resistance to autosomal dominant Alzheimer’s disease in an APOE3 Christchurch homozygote: a case report. Nat Med 25, 1680–1683. 10.1038/s41591-019-0611-3.

2. Hardy, J., and Selkoe, D.J. (2002). The amyloid hypothesis of Alzheimer’s disease: Progress and problems on the road to therapeutics. Science 297, 353–356.

3. Malpetti, M., Joie, R., and Rabinovici, G.D. (2022). Tau Beats Amyloid in Predicting Brain Atrophy in Alzheimer Disease: Implications for Prognosis and Clinical Trials. J Nucl Med 63, 830–832. 10.2967/jnumed.121.263694.

4. Lee, J.C., Kim, S.J., Hong, S., and Kim, Y. (2019). Diagnosis of Alzheimer’s disease utilizing amyloid and tau as fluid biomarkers. Exp Mol Med 51, 1–10. 10.1038/s12276-019-0250-2.

5. Bloom, G.S. (2014). Amyloid-beta and tau: the trigger and bullet in Alzheimer disease pathogenesis. JAMA Neurol 71, 505–508. 10.1001/jamaneurol.2013.5847.

6. Mahley, R.W., Weisgraber, K.H., and Huang, Y. (2006). Apolipoprotein E4: a causative factor and therapeutic target in neuropathology, including Alzheimer’s disease. Proc Natl Acad Sci U S A 103, 5644–5651.

7. Holtzman, D.M., Bales, K.R., Tenkova, T., Fagan, A.M., Parsadanian, M., Sartorius, L.J., Mackey, B., Olney, J., McKeel, D., Wozniak, D., and Paul, S.M. (2000). Apolipoprotein E isoform-dependent amyloid deposition and neuritic degeneration in a mouse model of Alzheimer’s disease. Proc Natl Acad Sci U S A 97, 2892–2897. 10.1073/pnas.050004797.

8. Lee, S., Devanney, N.A., Golden, L.R., Smith, C.T., Schwartz, J.L., Walsh, A.E., Clarke, H.A., Goulding, D.S., Allenger, E.J., Morillo-Segovia, G., et al. (2023). APOE modulates microglial immunometabolism in response to age, amyloid pathology, and inflammatory challenge. Cell Rep 42, 112196. 10.1016/j.celrep.2023.112196.

9. Wang, N., Wang, M., Jeevaratnam, S., Rosenberg, C., Ikezu, T.C., Shue, F., Doss, S.V., Alnobani, A., Martens, Y.A., Wren, M., et al. (2022). Opposing effects of apoE2 and apoE4 on microglial activation and lipid metabolism in response to demyelination. Mol Neurodegener 17, 75. 10.1186/s13024-022-00577-1.

10. Shi, Y., Yamada, K., Liddelow, S.A., Smith, S.T., Zhao, L., Luo, W., Tsai, R.M., Spina, S., Grinberg, L.T., Rojas, J.C., et al. (2017). ApoE4 markedly exacerbates tau-mediated neurodegeneration in a mouse model of tauopathy. Nature 549, 523–527. 10.1038/nature24016.

11. Liu, C.C., Murray, M.E., Li, X., Zhao, N., Wang, N., Heckman, M.G., Shue, F., Martens, Y., Li, Y., Raulin, A.C., et al. (2021). APOE3-Jacksonville (V236E) variant reduces self-aggregation and risk of dementia. Sci Transl Med 13, eabc9375. 10.1126/scitranslmed.abc9375.

12. De Strooper, B., Saftig, P., Craessaerts, K., Vanderstichele, H., Guhde, G., Annaert, W., Von Figura, K., and Van Leuven, F. (1998). Deficiency of presenilin-1 inhibits the normal cleavage of amyloid precursor protein. Nature 391, 387–390.

13. Sepulveda-Falla, D., Sanchez, J.S., Almeida, M.C., Boassa, D., Acosta-Uribe, J., Vila-Castelar, C., Ramirez-Gomez, L., Baena, A., Aguillon, D., Villalba-Moreno, N.D., et al. (2022). Distinct tau neuropathology and cellular profiles of an APOE3 Christchurch homozygote protected against autosomal dominant Alzheimer’s dementia. Acta Neuropathol. 10.1007/s00401-022-02467-8.

14. Nelson, M.R., Liu, P., Agrawal, A., Yip, O., Blumenfeld, J., Traglia, M., Kim, M.J., Koutsodendris, N., Rao, A., Grone, B., et al. (2023). The APOE-R136S mutation protects against APOE4-driven Tau pathology, neurodegeneration and neuroinflammation. Nat Neurosci 26, 2104–2121. 10.1038/s41593-023-01480-8.

15. Chen, Y., Song, S., Parhizkar, S., Lord, J., Zhu, Y., Strickland, M.R., Wang, C., Park, J., Tabor, G.T., Jiang, H., et al. (2023). APOE3ch alters microglial response and suppresses Abeta-induced tau seeding and spread. Cell. 10.1016/j.cell.2023.11.029.

16. Yoshiyama, Y., Higuchi, M., Zhang, B., Huang, S.M., Iwata, N., Saido, T.C., Maeda, J., Suhara, T., Trojanowski, J.Q., and Lee, V.M. (2007). Synapse loss and microglial activation precede tangles in a P301S tauopathy mouse model. Neuron 53, 337–351.

17. Keren-Shaul, H., Spinrad, A., Weiner, A., Matcovitch-Natan, O., Dvir-Szternfeld, R., Ulland, T.K., David, E., Baruch, K., Lara-Astaiso, D., Toth, B., et al. (2017). A Unique Microglia Type Associated with Restricting Development of Alzheimer’s Disease. Cell 169, 1276–1290 e1217. 10.1016/j.cell.2017.05.018.

18. Krasemann, S., Madore, C., Cialic, R., Baufeld, C., Calcagno, N., El Fatimy, R., Beckers, L., O’Loughlin, E., Xu, Y., Fanek, Z., et al. (2017). The TREM2-APOE Pathway Drives the Transcriptional Phenotype of Dysfunctional Microglia in Neurodegenerative Diseases. Immunity 47, 566–581 e569. 10.1016/j.immuni.2017.08.008.

19. Jicha, G.A., Bowser, R., Kazam, I.G., and Davies, P. (1997). Alz-50 and MC-1, a new monoclonal antibody raised to paired helical filaments, recognize conformational epitopes on recombinant tau. Journal of Neuroscience Research 48, 128–132. 10.1002/(SICI)1097-4547(19970415)48:2<128::AID-JNR5>3.0.CO;2-E.

20. Fu, H., Rodriguez, G.A., Herman, M., Emrani, S., Nahmani, E., Barrett, G., Figueroa, H.Y., Goldberg, E., Hussaini, S.A., and Duff, K.E. (2017). Tau Pathology Induces Excitatory Neuron Loss, Grid Cell Dysfunction, and Spatial Memory Deficits Reminiscent of Early Alzheimer’s Disease. Neuron 93, 533–541 e535. 10.1016/j.neuron.2016.12.023.

21. Busche, M.A., Wegmann, S., Dujardin, S., Commins, C., Schiantarelli, J., Klickstein, N., Kamath, T.V., Carlson, G.A., Nelken, I., and Hyman, B.T. (2019). Tau impairs neural circuits, dominating amyloid-beta effects, in Alzheimer models in vivo. Nat Neurosci 22, 57–64. 10.1038/s41593-018-0289-8.

22. Yuan, P., Zhang, M., Tong, L., Morse, T.M., McDougal, R.A., Ding, H., Chan, D., Cai, Y., and Grutzendler, J. (2022). PLD3 affects axonal spheroids and network defects in Alzheimer’s disease. Nature 612, 328–337. 10.1038/s41586-022-05491-6.

23. Nakamura, A., Cuesta, P., Fernandez, A., Arahata, Y., Iwata, K., Kuratsubo, I., Bundo, M., Hattori, H., Sakurai, T., Fukuda, K., et al. (2018). Electromagnetic signatures of the preclinical and prodromal stages of Alzheimer’s disease. Brain 141, 1470–1485. 10.1093/brain/awy044.

24. Bullitt, E. (1990). Expression of c-fos-like protein as a marker for neuronal activity following noxious stimulation in the rat. J Comp Neurol 296, 517–530. 10.1002/cne.902960402.

25. Deng, W., Aimone, J.B., and Gage, F.H. (2010). New neurons and new memories: how does adult hippocampal neurogenesis affect learning and memory? Nat Rev Neurosci 11, 339–350. 10.1038/nrn2822.

26. Vago, D.R., Bevan, A., and Kesner, R.P. (2007). The role of the direct perforant path input to the CA1 subregion of the dorsal hippocampus in memory retention and retrieval. Hippocampus 17, 977–987. 10.1002/hipo.20329.

27. Goettemoeller, A.M., Banks, E., Kumar, P., Olah, V.J., McCann, K.E., South, K., Ramelow, C.C., Eaton, A., Duong, D.M., Seyfried, N.T., et al. (2024). Entorhinal cortex vulnerability to human APP expression promotes hyperexcitability and tau pathology. Nat Commun 15, 7918. 10.1038/s41467-024-52297-3.

28. Parcellier, A., Tintignac, L.A., Zhuravleva, E., Cron, P., Schenk, S., Bozulic, L., and Hemmings, B.A. (2009). Carboxy-Terminal Modulator Protein (CTMP) is a mitochondrial protein that sensitizes cells to apoptosis. Cell Signal 21, 639–650. 10.1016/j.cellsig.2009.01.016.

29. Ono, H., Sakoda, H., Fujishiro, M., Anai, M., Kushiyama, A., Fukushima, Y., Katagiri, H., Ogihara, T., Oka, Y., Kamata, H., et al. (2007). Carboxy-terminal modulator protein induces Akt phosphorylation and activation, thereby enhancing antiapoptotic, glycogen synthetic, and glucose uptake pathways. Am J Physiol Cell Physiol 293, C1576–1585. 10.1152/ajpcell.00570.2006.

30. Sayed, F.A., Kodama, L., Fan, L., Carling, G.K., Udeochu, J.C., Le, D., Li, Q., Zhou, L., Wong, M.Y., Horowitz, R., et al. (2021). AD-linked R47H-TREM2 mutation induces disease-enhancing microglial states via AKT hyperactivation. Sci Transl Med 13, eabe3947. 10.1126/scitranslmed.abe3947.

31. de Lima, I.B.Q., Cardozo, P.L., Fahel, J.S., Lacerda, J.P.S., Miranda, A.S., Teixeira, A.L., and Ribeiro, F.M. (2023). Blockade of mGluR5 in astrocytes derived from human iPSCs modulates astrocytic function and increases phagocytosis. Front Immunol 14, 1283331. 10.3389/fimmu.2023.1283331.

32. Balcar, V.J., Zeman, T., Janout, V., Janoutová, J., Lochman, J., and Šerý, O. (2021). Single Nucleotide Polymorphism rs11136000 of CLU Gene (Clusterin, ApoJ) and the Risk of Late-Onset Alzheimer’s Disease in a Central European Population. Neurochem Res 46, 411–422. 10.1007/s11064-020-03176-y.

33. Chen, F., Swartzlander, D.B., Ghosh, A., Fryer, J.D., Wang, B., and Zheng, H. (2021). Clusterin secreted from astrocyte promotes excitatory synaptic transmission and ameliorates Alzheimer’s disease neuropathology. Mol Neurodegener 16, 5. 10.1186/s13024-021-00426-7.

34. Yang, Y., Gozen, O., Watkins, A., Lorenzini, I., Lepore, A., Gao, Y., Vidensky, S., Brennan, J., Poulsen, D., Won Park, J., et al. (2009). Presynaptic regulation of astroglial excitatory neurotransmitter transporter GLT1. Neuron 61, 880–894. 10.1016/j.neuron.2009.02.010.

35. Jiang, W., Yang, W., Yang, W., Zhang, J., Pang, D., Gan, L., Luo, L., Fan, Y., Liu, Y., and Chen, M. (2013). Identification of Tmem10 as a novel late-stage oligodendrocytes marker for detecting hypomyelination. Int J Biol Sci 10, 33–42. 10.7150/ijbs.7526.

36. Zhu, Q., Tan, Z., Zhao, S., Huang, H., Zhao, X., Hu, X., Zhang, Y., Shields, C.B., Uetani, N., and Qiu, M. (2015). Developmental expression and function analysis of protein tyrosine phosphatase receptor type D in oligodendrocyte myelination. Neuroscience 308, 106–114. 10.1016/j.neuroscience.2015.08.062.

37. Giussani, P., Prinetti, A., and Tringali, C. (2021). The role of Sphingolipids in myelination and myelin stability and their involvement in childhood and adult demyelinating disorders. J Neurochem 156, 403–414. 10.1111/jnc.15133.

38. Boland, S., Swarup, S., Ambaw, Y.A., Malia, P.C., Richards, R.C., Fischer, A.W., Singh, S., Aggarwal, G., Spina, S., Nana, A.L., et al. (2022). Deficiency of the frontotemporal dementia gene GRN results in gangliosidosis. Nat Commun 13, 5924. 10.1038/s41467-022-33500-9.

39. Fitz, N.F., Nam, K.N., Wolfe, C.M., Letronne, F., Playso, B.E., Iordanova, B.E., Kozai, T.D.Y., Biedrzycki, R.J., Kagan, V.E., Tyurina, Y.Y., et al. (2021). Phospholipids of APOE lipoproteins activate microglia in an isoform-specific manner in preclinical models of Alzheimer’s disease. Nat Commun 12, 3416. 10.1038/s41467-021-23762-0.

40. Wang, C., Fan, L., Khawaja, R.R., Liu, B., Zhan, L., Kodama, L., Chin, M., Li, Y., Le, D., Zhou, Y., et al. (2022). Microglial NF-κB drives tau spreading and toxicity in a mouse model of tauopathy. Nat Commun 13, 1969. 10.1038/s41467-022-29552-6.

41. Udeochu, J.C., Amin, S., Huang, Y., Fan, L., Torres, E.R.S., Carling, G.K., Liu, B., McGurran, H., Coronas-Samano, G., Kauwe, G., et al. (2023). Tau activation of microglial cGAS-IFN reduces MEF2C-mediated cognitive resilience. Nat Neurosci 26, 737–750. 10.1038/s41593-023-01315-6.

42. Sun, G.G., Wang, C., Mazzarino, R.C., Perez-Corredor, P.A., Davtyan, H., Blurton-Jones, M., Lopera, F., Arboleda-Velasquez, J.F., and Shi, Y. (2024). Microglial APOE3 Christchurch protects neurons from Tau pathology in a human iPSC-based model of Alzheimer’s disease. Cell Rep 43, 114982. 10.1016/j.celrep.2024.114982.

43. McQuade, A., Coburn, M., Tu, C.H., Hasselmann, J., Davtyan, H., and Blurton-Jones, M. (2018). Development and validation of a simplified method to generate human microglia from pluripotent stem cells. Mol Neurodegener 13, 67. 10.1186/s13024-018-0297-x.

44. Lama, L., Adura, C., Xie, W., Tomita, D., Kamei, T., Kuryavyi, V., Gogakos, T., Steinberg, J.I., Miller, M., Ramos-Espiritu, L., et al. (2019). Development of human cGAS-specific small-molecule inhibitors for repression of dsDNA-triggered interferon expression. Nat Commun 10, 2261. 10.1038/s41467-019-08620-4.

45. Lopez-Lee, C., Kodama, L., Fan, L., Zhu, D., Zhu, J., Wong, M.Y., Ye, P., Norman, K., Foxe, N.R., Ijaz, L., et al. (2024). Tlr7 drives sex differences in age- and Alzheimer’s disease-related demyelination. Science 386, eadk7844. 10.1126/science.adk7844.

46. Palop, J.J., and Mucke, L. (2010). Amyloid-beta-induced neuronal dysfunction in Alzheimer’s disease: from synapses toward neural networks. Nat Neurosci 13, 812–818. nn.2583 [pii]10.1038/nn.2583 [doi].

47. Busche, M.A., and Konnerth, A. (2015). Neuronal hyperactivity--A key defect in Alzheimer’s disease? Bioessays 37, 624–632. 10.1002/bies.201500004.

48. Verret, L., Mann, E.O., Hang, G.B., Barth, A.M., Cobos, I., Ho, K., Devidze, N., Masliah, E., Kreitzer, A.C., Mody, I., et al. (2012). Inhibitory interneuron deficit links altered network activity and cognitive dysfunction in Alzheimer model. Cell 149, 708–721. 10.1016/j.cell.2012.02.046.

49. Hijazi, S., Heistek, T.S., Scheltens, P., Neumann, U., Shimshek, D.R., Mansvelder, H.D., Smit, A.B., and van Kesteren, R.E. (2020). Early restoration of parvalbumin interneuron activity prevents memory loss and network hyperexcitability in a mouse model of Alzheimer’s disease. Mol Psychiatry 25, 3380–3398. 10.1038/s41380-019-0483-4.

50. Algamal, M., Russ, A.N., Miller, M.R., Hou, S.S., Maci, M., Munting, L.P., Zhao, Q., Gerashchenko, D., Bacskai, B.J., and Kastanenka, K.V. (2022). Reduced excitatory neuron activity and interneuron-type-specific deficits in a mouse model of Alzheimer’s disease. Commun Biol 5, 1323. 10.1038/s42003-022-04268-x.

51. Xu, Y., Zhao, M., Han, Y., and Zhang, H. (2020). GABAergic Inhibitory Interneuron Deficits in Alzheimer’s Disease: Implications for Treatment. Front Neurosci 14, 660. 10.3389/fnins.2020.00660.

52. Iaccarino, H.F., Singer, A.C., Martorell, A.J., Rudenko, A., Gao, F., Gillingham, T.Z., Mathys, H., Seo, J., Kritskiy, O., Abdurrob, F., et al. (2016). Gamma frequency entrainment attenuates amyloid load and modifies microglia. Nature 540, 230–235. 10.1038/nature20587.

53. Li, G., Bien-Ly, N., Andrews-Zwilling, Y., Xu, Q., Bernardo, A., Ring, K., Halabisky, B., Deng, C., Mahley, R.W., and Huang, Y. (2009). GABAergic interneuron dysfunction impairs hippocampal neurogenesis in adult apolipoprotein E4 knockin mice. Cell Stem Cell 5, 634–645. 10.1016/j.stem.2009.10.015.

54. Pineda, J.R., and Encinas, J.M. (2016). The Contradictory Effects of Neuronal Hyperexcitation on Adult Hippocampal Neurogenesis. Front Neurosci 10, 74. 10.3389/fnins.2016.00074.

55. Sharma, A., Kazim, S.F., Larson, C.S., Ramakrishnan, A., Gray, J.D., McEwen, B.S., Rosenberg, P.A., Shen, L., and Pereira, A.C. (2019). Divergent roles of astrocytic versus neuronal EAAT2 deficiency on cognition and overlap with aging and Alzheimer’s molecular signatures. Proc Natl Acad Sci U S A 116, 21800–21811. 10.1073/pnas.1903566116.

56. Masliah, E., Alford, M., DeTeresa, R., Mallory, M., and Hansen, L. (1996). Deficient glutamate transport is associated with neurodegeneration in Alzheimer’s disease. Ann Neurol 40, 759–766. 10.1002/ana.410400512.

57. Kong, Q., Chang, L.C., Takahashi, K., Liu, Q., Schulte, D.A., Lai, L., Ibabao, B., Lin, Y., Stouffer, N., Das Mukhopadhyay, C., et al. (2014). Small-molecule activator of glutamate transporter EAAT2 translation provides neuroprotection. J Clin Invest 124, 1255–1267. 10.1172/JCI66163.

58. Blumenreich, S., Nehushtan, T., Barav, O.B., Saville, J.T., Dingjan, T., Hardy, J., Fuller, M., and Futerman, A.H. (2022). Elevation of gangliosides in four brain regions from Parkinson’s disease patients with a GBA mutation. NPJ Parkinsons Dis 8, 99. 10.1038/s41531-022-00363-2.

59. Asai, H., Ikezu, S., Tsunoda, S., Medalla, M., Luebke, J., Haydar, T., Wolozin, B., Butovsky, O., Kugler, S., and Ikezu, T. (2015). Depletion of microglia and inhibition of exosome synthesis halt tau propagation. Nat Neurosci 18, 1584–1593. 10.1038/nn.4132.

60. Shi, Y., Yamada, K., Liddelow, S.A., Smith, S.T., Zhao, L., Luo, W., Tsai, R.M., Spina, S., Grinberg, L.T., Rojas, J.C., et al. (2017). ApoE4 markedly exacerbates tau-mediated neurodegeneration in a mouse model of tauopathy. Nature 549, 523–527. 10.1038/nature24016.

61. Chen, X., Firulyova, M., Manis, M., Herz, J., Smirnov, I., Aladyeva, E., Wang, C., Bao, X., Finn, M.B., Hu, H., et al. (2023). Microglia-mediated T cell infiltration drives neurodegeneration in tauopathy. Nature 615, 668–677. 10.1038/s41586-023-05788-0.

62. Marino, C., Perez-Corredor, P., O’Hare, M., Heuer, A., Chmielewska, N., Gordon, H., Chandrahas, A.S., Gonzalez-Buendia, L., Delgado-Tirado, S., Doan, T.H., et al. (2024). APOE Christchurch-mimetic therapeutic antibody reduces APOE-mediated toxicity and tau phosphorylation. Alzheimers Dement 20, 819–836. 10.1002/alz.13436.

63. Gordts, P., Foley, E.M., Lawrence, R., Sinha, R., Lameda-Diaz, C., Deng, L., Nock, R., Glass, C.K., Erbilgin, A., Lusis, A.J., et al. (2014). Reducing macrophage proteoglycan sulfation increases atherosclerosis and obesity through enhanced type I interferon signaling. Cell Metab 20, 813–826. 10.1016/j.cmet.2014.09.016.

64. Siegle, J.H., López, A.C., Patel, Y.A., Abramov, K., Ohayon, S., and Voigts, J. (2017). Open Ephys: an open-source, plugin-based platform for multichannel electrophysiology. Journal of Neural Engineering 14, 045003. 10.1088/1741-2552/aa5eea.

65. Lopes, G., and Monteiro, P. (2021). New Open-Source Tools: Using Bonsai for Behavioral Tracking and Closed-Loop Experiments. Frontiers in Behavioral Neuroscience 15. 10.3389/fnbeh.2021.647640.

66. Mathis, A., Mamidanna, P., Cury, K.M., Abe, T., Murthy, V.N., Mathis, M.W., and Bethge, M. (2018). DeepLabCut: markerless pose estimation of user-defined body parts with deep learning. Nature Neuroscience 21, 1281–1289. 10.1038/s41593-018-0209-y.

67. Chi, J., Crane, A., Wu, Z., and Cohen, P. (2018). Adipo-Clear: A Tissue Clearing Method for Three-Dimensional Imaging of Adipose Tissue. J Vis Exp. 10.3791/58271.

68. Hou, Y., Zhang, Q., Liu, H., Wu, J., Shi, Y., Qi, Y., Shao, M., Yang, Z., Lu, J., Wu, Z., et al. (2021). Topographical organization of mammillary neurogenesis and efferent projections in the mouse brain. Cell Rep 34, 108712. 10.1016/j.celrep.2021.108712.

69. Kim, Y., Yang, G.R., Pradhan, K., Venkataraju, K.U., Bota, M., Garcia Del Molino, L.C., Fitzgerald, G., Ram, K., He, M., Levine, J.M., et al. (2017). Brain-wide Maps Reveal Stereotyped Cell-Type-Based Cortical Architecture and Subcortical Sexual Dimorphism. Cell 171, 456–469 e422. 10.1016/j.cell.2017.09.020.

70. Muñoz-Castañeda, R., Palaniswamy, R., Palmer, J., Drewes, R., Elowsky, C., Hirokawa, K.E., Cain, N., Venkataraju, K.U., Harris, J.A., and Osten, P. (2024). Brain Structural Organization Revealed by Unbiased Cell-Type Distribution Clustering. bioRxiv, 2024.2010.2002.615922. 10.1101/2024.10.02.615922.

71. Klein, S., Staring, M., Murphy, K., Viergever, M.A., and Pluim, J.P. (2010). elastix: a toolbox for intensity-based medical image registration. IEEE Trans Med Imaging 29, 196–205. 10.1109/TMI.2009.2035616.

72. Folch, J., Lees, M., and Sloane Stanley, G.H. (1957). A simple method for the isolation and purification of total lipides from animal tissues. J Biol Chem 226, 497–509.

73. Anders, S., Pyl, P.T., and Huber, W. (2015). HTSeq--a Python framework to work with high-throughput sequencing data. Bioinformatics 31, 166–169. 10.1093/bioinformatics/btu638.

